# Stromal mediated DNA damage promotes high grade serous ovarian cancer initiation

**DOI:** 10.1101/2024.05.23.595550

**Authors:** Geyon L. Garcia, Taylor Orellana, Grace Gorecki, Leonard G. Frisbie, Roja Baruwal, Ester Goldfield, Ian Beddows, Ian P. MacFawn, Ananya K. Britt, Macy M. Hale, Hui Shen, Ronald Buckanovich, Toren Finkel, Ronny Drapkin, T. Rinda Soong, Tullia C. Bruno, Huda I. Atiya, Lan Coffman

## Abstract

The fundamental steps in high-grade serous ovarian cancer (HGSOC) initiation are unclear, thus providing critical barriers to the development of prevention or early detection strategies for this deadly disease. Increasing evidence demonstrates most HGSOC starts in the fallopian tube epithelium (FTE). Current models propose HGSOC initiates when FTE cells acquire increasing numbers of mutations allowing cells to evolve into serous tubal intraepithelial carcinoma (STIC) precursors and then to full blown cancer. Here we report that epigenetically altered mesenchymal stem cells (termed high risk MSC-hrMSCs) can be detected prior to the formation of ovarian cancer precursor lesions. These hrMSCs drive DNA damage in the form of DNA double strand breaks in FTE cells while also promoting the survival of FTE cells in the face of DNA damage. Indicating the hrMSC may actually drive cancer initiation, we find hrMSCs induce full malignant transformation of otherwise healthy, primary FTE resulting in metastatic cancer *in vivo*. Further supporting a role for hrMSCs in cancer initiation in humans, we demonstrate that hrMSCs are highly enriched in BRCA1/2 mutation carriers and increase with age. Combined these findings indicate that hrMSCs may incite ovarian cancer initiation. These findings have important implications for ovarian cancer detection and prevention.

## Introduction

Ovarian cancer is the most lethal gynecologic cancer, with over 13,000 deaths predicted to occur in 2023^1^. High-grade serous ovarian carcinoma (HGSOC) is the most common ovarian cancer subtype, with more than 70% of patients presenting with metastatic disease at the time of diagnosis^2^. This is attributed to multiple factors, including a non-specific and gradually worsening symptom burden and effective screening or early detection strategies. There is a critical need to identify and understand the mechanisms of HGSOC initiation, which can then be leveraged to derive effective prevention and early diagnosis approaches.

Due to the inability to screen for ovarian cancer, the current approach to primary prevention is surgical castration with bilateral salpingo-oophorectomies in women with BRCA1 or −2 germline mutations. BRCA1 and BRCA2 mutations are well-characterized risk factors conveying a roughly 39-58% or 13-29% lifetime risk of developing HGSOC, respectively ^2^. However, the majority of HGSOC tumors develop in the absence of germline BRCA1 or −2 mutations, suggesting that malignant transformation is achieved by other, currently undefined, mechanisms. Convincing evidence has demonstrated that the majority of HGSOC tumors arise from cells in the fallopian tube epithelium (FTE), with the immediate precursor lesion of HGSOC termed a serous tubal intraepithelial carcinoma (STIC)^3–6^. STIC lesions are a series of *p53*-mutated epithelial cells demonstrating atypical morphology and increased proliferative index indicative of increased malignant behavior ^7,89^. Research on STIC lesions has largely focused on changes within epithelial cells, while the microenvironment surrounding STIC lesions has largely been unexplored. It is clear the microenvironment plays a critical role in the pathophysiology of HGSOC; however, its role in tumor initiation is undefined. Evidence from other cancer types demonstrates that changes in the stromal microenvironment may create a permissive or even causative environment that promotes cancer initiation^10–12^. The role of the fallopian tube microenvironment in HGSOC initiation is a critical gap in our current understanding of this disease.

Our group previously demonstrated that mesenchymal stromal/stem cells (MSCs), a multipotent stromal progenitor cell found in most adult tissues, can be reprogramed by cancer cells to form cancer-associated mesenchymal stem cells (CA-MSCs)^13^. These HGSOC-educated CA-MSCs exhibit a tumor-supportive phenotype that enhances tumor cell growth, confers chemoresistance, enriches the cancer stem-like cell pool, promotes angiogenesis, and aids in the metastatic spread of HGSOC tumor cells^13^. The pro-tumorigenic phenotype of CA-MSCs is accompanied by a unique DNA methylation, RNA expression, and protein expression profile that differs from non-tumorigenic or normal MSCs (nMSCs). Using these data, we discovered that CA-MSCs rely on expression of the transcription factor Wilms Tumor Protein 1 (WT1) to carry out their pro-tumorigenic function^13^.

Here, we identified a subset of fallopian tube-derived MSCs from women without cancer that phenocopy cancer-educated CA-MSCs. This group of MSCs exhibited similar epigenetic and transcriptomic changes to CA-MSCs, including high expression of WT1. This subset of MSCs is enriched in women with BRCA1 or −2 mutations, increases with age, and is enriched within the stroma surrounding STIC lesions. Additionally, MSCs exhibit a tumor-supportive phenotype like their cancer-associated counterparts. That is, these MSCs, which we refer to as high-risk MSCs (hrMSCs), enhance tumor cell growth, chemoresistance, and sphere formation and increase the proliferation and “stemness” of FTE cells. Most notably, hrMSCs trigger full malignant transformation, *in vivo*, of primary, non-cancerous FTE cells, thus indicating that hrMSCs are a mediator of HGSOC initiation. In support of these findings, we demonstrate that hrMSCs are potent inducers of DNA damage in FTE and promote recovery of FTE following oxidative stress; two mechanisms that are linked to HGSOC initiation. This work describes the tumor-initiating function of hrMSCs and begins to elucidate an important mechanism of HGSOC formation.

## Methods

### nMSCs/hrMSCs

Mesenchymal stem cells (MSCs) were isolated from fallopian tubes of women with or without BRCA mutations undergoing gynecologic surgery for benign indications or risk-reducing salpingo-oophorectomies without findings of cancer or pre-cancerous lesions upon standard pathologic examination. Using previously described protocols, fresh fallopian tubes were cut into < 1mm^3^ pieces and plated in 6-well-plates ^13^. Following ISCT guidelines on the minimal criteria for defining multipotent mesenchymal stem cells, MSCs were selected for plastic adherence and cell surface marker expression (CD105+, CD90+, CD73+, CD45-, CD34-, CD14-, CD19-)^14^. Adipocyte, osteocyte, and chondrocyte differentiation capacity was also verified. MSCs were propagated using Mammary Epithelial cell Basal Medium (MEBM) supplemented with 10% heat-inactivated FBS (fetal bovine serum), 1X B27, 20 ng/mL EGF, 1 ng/mL hydrocortisone, 5 μg/mL insulin, 100 μM β-mercaptoethanol, 10 ng/mL β-FGF, 1% Penicillin/Streptomycin, and 20 μg/mL gentamicin. MSCs were used for functional experiments for, at most, 8-10 passages.

### Fallopian tube epithelium (FTE)

Using fresh normal human fallopian tube tissue, FTEs were isolated following a modified version of a previously described protocol^15^. Briefly, fresh tissue was mechanically and enzymatically dissociated into a single-cell suspension, plated, and passaged on tissue culture-treated placental collagen IV (Sigma Aldrich; C7521)-coated plates. The single-cell suspension was then cultured in Dulbecco’s modification of Eagle’s medium/Ham’s F-12 50/50 mix (DMEM-Ham’s F12) supplemented with 2% Ultroser G serum substitute and 1% penicillin/streptomycin. FTE cells were validated for expression of panCK and negative for MSC markers CD105, CD90 and CD73. Immortalized FT190 control cells were utilized in parallel where possible. For all adherent and non-adherent co-culture experiments, FTE cells were plated at a 1:1 ratio with MSCs. This ratio was chosen because FTE rapidly undergo replicative senescence in *ex vivo* culture. Higher ratios of FTE to MSCs resulted in poor FTE growth kinetics.

### Methylation data processing

EPIC array IDATs from GSE138072 were combined with an additional 12 EPIC array samples for a dataset of 14 FT, 23 OM, and 7 OV samples (N=44 total samples). These samples were further differentiated as nMSCs (N=20), CA-MSCs (N=14), and hrMSCs with heterozygous BRCA mutations (N=10). IDATs were processed using sesame v1.8.2 with the openSesame function and default parameters (Zhou et al., 2018). Array probes were excluded if they had more than 10% missing data across all samples; this resulted in exclusion of 474,808 probes.

### Probe selection

The top 1,000 probes with largest average increase in methylation in CA-MSCs relative to normal MSCs, as well as the 1,000 with the largest average decrease in methylation in CA-MSCs, were selected as these probes best discriminate the normal and cancer-associated MSCs. hrMSCs were not included in this probe selection. All samples were included in the selection of the top variable probes, which were selected by ranking the standard deviation across all samples.

UMAP analysis was done using the uwot package v0.1.14 (McInnes et al., 2018). The umap was done using the top discriminating probes for CA-MSC–MSC scaled with column mean of 0 and variance of 1. The umap function was run with default parameters, except the spread option was set to 2, and the effective minimum distance between embedded points was set to 0.001. Heatmaps were generated using ComplexHeatmap v2.14.0^16,1716^.

### FTE proliferation assay

FTE cells were stained with CellTrace Violet (CTV - Invitrogen <u>C34571</u>) following the manufacturer’s protocol. 5 × 10^3^ CTV-labeled FT cells were cultured alone, with 5 × 10^3^ nMSCs or with 5 × 10^3^ hrMSCs in a 12-well plate. The FT cell numbers were followed for 4 days. Cells were collected from two wells per condition per day and counted using a hemocytometer. Flow cytometric analysis was used to determine the percentage of the CTV-labeled FT to quantify viable FTs per condition. Total FT cells (CTV-labeled) = total cell number × CTV positive cell %.

### Sphere formation assay

Primary FTE cells were stained with CTV, following the manufacturer’s protocol. Using ultra-low-adherence 96-well plates, labelled FTEs were cultured alone (1×10^3^ cells/well), with hrMSCs (1×10^3^ total cells/well), or with nMSCs (1×10^3^ total cells/well) in 300 µl MEBM media supplemented with 1X B27, 20 ng/mL EGF, 1 ng/mL hydrocortisone, 5 μg/mL insulin, 100 μM β-mercaptoethanol, 10 ng/mL β-FGF, 1% penicillin/streptomycin, and 20 μg/mL gentamicin. Spheres were counted in the entire well at 7 days from plating (> 4 cells spheroid, at least 1 FTE per sphere).

### Cell adhesion assay

3×10^4 nMSCs, hrMScs, or CA-MSCs were cultured overnight in a 12-well plate to form a monolayer. 3×10^4^ HGSOC cells (OVSAHO, OVCAR3, or pt412) were stained with CTV and then added to the cultured nMSCs, hrMScs, or CA-MSCs. After 30 minutes, cells were then washed twice with PBS, and the attached CTV-labeled HGSOC cells were counted using a fluorescence microscope. Experiments were repeated independently three times per tumor cell type.

### Quantitative real-time PCR

Using RNeasy Mini Kit and on-column DNase treatment (Qiagen, 28106), RNA was isolated from MSCs. Samples were then processed for cDNA synthesis using SuperScript III First Strand Synthesis System (random hexamer; Invitrogen, 18080–051). SYBR-green based RT-qPCR (Applied Biosystems, 4472908) was performed using CFX96 Real-Time System, with GAPDH as the reference gene. Samples were run for 40 cycles.

### Flow cytometric analysis

For quantification of WT1 expression, MSCs were washed with PBS, trypsonized into a single-cell suspension, and counted. MSCs were processed using the intracellular fixation and permeabilization buffer set (eBioscience; 88-8824-00). Briefly, 1×10^6^ cells were resuspended in IC fixation buffer and fixed at room temperature (RT) for 20 minutes. Cells were washed twice with IC permeabilization buffer and subsequently stained with mouse anti-human WT1 405 (R & D; IC57291V; 1:100) for 20 minutes at RT, protected from light. MSCs were washed twice with IC permeabilization buffer, resuspended in PBS, and analyzed using the CytoFLEX 4L cytometer. Ovarian cancer cell lines (e.g., OVCAR3) and primary CA-MSC cell lines were used as positive controls^13,18–22^.

For quantification of the CellROX (Life Technologies; C10448) oxidation probe, minor adjustments were made to the manufacturer’s protocol. Briefly, MSCs were treated with complete media +/- 50-100 µM hydrogen peroxide and 1-10µM Trolox for 30 minutes. Afterwards, MSCs were rinsed with complete media and stained with 5 µM of the CellROX Deep Red (DR) probe for 30 minutes at 37°C for 30 minutes. MSCs were rinsed three times with PBS and processed using the IC Flow kit as described above. MSCs were run on the cytometer within the suggested 2-hour interval. To correlate WT1 expression with CellROX DR, MSCs were stained with WT1 conjugates for 20 minutes after permeabilization.

For quantification of 4-hydroxynonenal (4-HNE), we relied on commercially available 4-HNE antibody conjugates. Briefly, MSC controls were pretreated with either 50-100 µM hydrogen peroxide or 5-10 µM Trolox for 30 minutes. Afterwards, MSCs and their controls were rinsed with complete media and processed using the IC flow kit. After permeabilization, MSCs were stained with mouse anti-4-HNE (Invitrogen; 12F7; MA5-45792; 1:100) or with mouse anti-MDA (Invitrogen; 6H6; MA5-45801; 1:100) conjugated antibodies for 20 minutes at RT. Cells were rinsed with 1X PBS three times and analyzed using the CytoFLEX 4L cytometer.

### Vectra multispectral imaging and analysis

For the multispectral immunofluorescence experiments, Akoya Bioscience’s Vectra MOTIF imaging pipeline and reagents were used. Auto staining of the panels was carried out on a Leica Bond Rx stainer, and the resulting stained slides were scanned using a Vectra Polaris imager. Subsequently, digital image files in QPTIFF format were downloaded and unmixed using Akoya Biosciences InForm software. Qupath software was utilized to identify Regions of Interest (ROIs) for phenotype analysis.

To quantify nMSC and hrMSC populations from our samples, we used QuPath v0.3.2 image analysis software. MOTIF QPTiffs were loaded into QuPath. Cell segmentation was performed based on nuclei (DAPI). We created a cell classifier, detecting cells that were positive or negative for the antibodies of interest (CD90, CD45, CD73, CD105, WT1, PanCK). The classifier was used to generate a phenotype algorithm applied to our cohort. We established that nMSCs had to be negative for CD45 and PanCK and positive for three MSC markers (CD90, 73, 105) and negative for WT1, while CA-MSCs were negative for CD45 and PanCK, positive for the same three MSC markers (CD90, 73, 105) and positive for WT1. We quantified the nMSCs and hrMSCs, normalized to the area of the analyzed ROI.

### DNA damage assay

NMSCs and hrMSCs were passaged until they were 40-60% confluent. For conditioned media experiments, MSC medium was replaced with complete MSC medium 24 hours prior to FTE treatment. At this time, 2,500 primary FTE or FT190 control cells were seeded in chamber slides. FTE cells were allowed to adhere overnight. Twenty-four hours later, MSC-conditioned medium was spun down at 4°C, 1,500 rpm, for 5 minutes to remove cells and cellular debris. FTE medium was aspirated off, and 500 µL of conditioned medium was added to cells. FTE cells were incubated in conditioned medium for up to 4 hours, then assayed for γH2AX (EMD Millipore; 05-636-1) fluorescence intensity and 53BP1 (Abcam; ab175933) foci by immunofluorescence. For co-culture experiments, 40-60% confluent MSC cultures were dissociated and seeded with FTE or FT190 control cells at a 1:1 ratio and allowed to adhere overnight. Twenty-four hours after co-culture, cells were assayed for γH2AX intensity and 53BP1 foci by immunofluorescence.

### FTE recovery assays

1,250-5,000 patient-derived primary FTE cells were plated in 96-well plates. Twenty-four hours later, cells were treated with 50 mM H_2_O_2_ for 10 minutes at RT. Cells were rinsed with PBS once and supplemented with conditioned medium. Conditioned medium was taken from 40-60% confluent nMSC or hrMSC cultures and was spun down at 4°C, 1,500 rpm, for 5 minutes to remove cells and cellular debris. nMSCs and hrMSCs from the same patient were matched for each experiment. Every 2 days, FTE medium was aspirated and replaced with 100 µL of conditioned medium. 20 µL of room-temperature MTS reagent (Celltiter96 AQ_ueous_ One Solution; Promega; G3582) was added to each well, using a multichannel pipette. Cells were incubated with the MTS reagent at 37°C for 4 hours. Absorbance was read on a plate reader at 490 nm.

### Immunofluorescence/immunohistochemistry staining

Cells were seeded in chamber slides and harvested at 24 hours. For γH2AX staining, cells were pre-extracted for 1 minute with pre-extraction buffer (0.6M EGTA, 0.5M PIPES, 0.5M MgSO_4_, 3.0M KCl, 1% Triton-X), then fixed for 20 minutes with ice cold 4% PFA in PBS. Pre-extraction was omitted for 53BP1 foci and MDA/4-HNE experiments. Cells were washed twice with PBS and permeabilized with 0.5% Triton-X in PBS for 10 minutes at RT. MDA/4HNE experiments were permeabilized with 0.1% Triton-X in PBS. Cells were washed twice with PBS, then blocked for 1 hour at RT with 100% SuperBlock (Thermo Scientific; 37535). Primary antibodies for DNA damage assays were diluted in PBS with 10% SuperBlock at the following dilutions: anti-human γH2AX Ser139 (EMD Millipore; JBW301; 1:200), goat anti-mouse IgG AF546 (Invitrogen; A11030; 1:2000), anti-human 53BP1 (Abcam; EPR2172; 1:200), goat anti-rabbit IgG AF488 (Abcam; ab150077; 1:4000) or donkey anti-rabbit AF647 (Abcam; ab150063; 1:4000). Cells were stained for 1 hour at RT then washed six times with PBS. Secondary antibodies were diluted as above and incubated on cells for 1 hour at RT. Cells were washed six times with PBS, mounted with Prolong Diamond Antifade Mountant with DAPI (Invitrogen; P36962) and allowed to dry overnight, protected from light.

For experiments utilizing the CellROX Green stain, cells were initially stained with live-cell F-actin (Spirochrome; SC001) for 1 hour at 37°C. Cells were washed three times with PBS and stained with 5 µM CellROX Green in PBS for 30 minutes. Cells were processed per manufacturer’s recommendations. Briefly, cells were fixed in IC fixation buffer and mounted using Prolong Diamond Antifade Mountant with DAPI. MSC controls were pretreated with either 50-100 µM hydrogen peroxide or 5-10 µM Trolox for 1 hour. Cells were rinsed three times with PBS and subsequently fixed with 4% PFA for 20 minutes at RT. Afterwards, MSCs and their controls were stained with F-actin, rinsed three times with PBS, and mounted with Prolong Diamond Antifade Mountant with DAPI. Slides were imaged on a Leica Thunder DMi8 fluorescent microscope within 24 hours of staining.

#### Image Quantification

All slides were imaged on a Leica Thunder DMi8 fluorescent microscope. All images were deconvolved prior to quantification in ImageJ. For image quantification, three or four random fields were taken, using the 20X objective for each condition. Fluorescence intensity (integrated density) for nuclear γH2AX, cytoplasmic 4-HNE and cytoplasmic MDA were quantified using ImageJ. 53BP1 foci per nuclei were also quantified using ImageJ.

### Lentiviral transduction

Lentiviral particles endcoding *PRKAA1*-mGFP (RC218572L2V) or empty vector mGFP control constructs (PS100071V) were purchased from OriGene. Briefly, 3×10^4^ hrMSCs were transduced with a multiplicity of infection (MOI) of 10 viral particles per cell. Empty vector viral particles at an MOI of 10 were used as negative controls. Virus medium was removed 24 hours after transduction, and cells were allowed to recover until they were 70% confluent (1-2 days). Cells were passaged and sorted on mGFP. GFP+ cells were validated for AMPKα1 overexpression and used immediately for experiments.

### Western blotting

Western blot was used to quantify the amount of total and phosphorylated (T172) AMPKα1. MSCs were rinsed with 1X PBS and centrifuged at 1,500 rpm for 5 minutes at 4°C. Pellets were lysed with RIPA (Thermo Scientific; 89900) containing 1X phosphatase inhibitor (Roche; 04906837001) and 1X protease inhibitor (Roche; 11697498001) for 30 minutes on ice. Lysates were centrifuged at 13,000xG for 5 minutes at 4°C. Supernatants were collected, and protein content was determined by BCA (Thermo Scientific; 23227). 30 µM of protein per sample was linearized with 1X LDS sample buffer (Invitrogen; NP0007) and loaded into 4-12% Bis-Tris precast gels (Invitrogen; NW04125). Samples were electrophoresed for 1.5 hours at 115 V and transferred onto nitrocellulose membranes (Cytiva; 10600001) using the Thermofisher semi-wet transfer apparatus. Membranes were blocked for 1 hour at RT, using TBS Licor Buffer (927-60001) with 0.01% Tween 20, then stained using rabbit anti-human pAMPKα1 (Cell Signalling; 40H9; 2535S; 1:500), rabbit anti-human AMPKα1 (Cell Signalling; 2535S; 1:500), and rabbit anti-human β-actin (Abcam; ab8227; 1:2000) overnight, rocking, at 4°C. Membranes were washed three times with 0.01% TBS-T and stained with goat anti-rabbit IR Dye 800CW (Licor; 926-32211; 1:10,000) or donkey anti-mouse IR Dye 680RD (Licor; 926-68072; 1:10,000) for 1 hour at RT. Membranes were washed three times with 0.01% TBS-T and imaged using an Odyssey CLx fluorescence imaging system. Band densitometry was determined in ImageJ.

### Mouse model tumor initiation

Experimental procedures were performed in accordance with the protocol approved by the Institutional Animal Care and Use Committee at the University of Pittsburgh. 6-to 8-week-old female NSG (NOD scid gamma) mice were used to assess tumor initiation capacity. An equivalent number of long-term organoids of *p53*-mutated FTE cells alone, with hrMSCs, or with nMSCs were injected into the inguinal mammary fat pad of NSG mice to allow to us to palpate tumors. Mice were monitored by palpating the injection site for tumor initiation. For secondary initiation studies, the primary *p53^mut^* tumors and two metastatic tumors were grown ex vivo. Mouse cells were depleted (see Mouse cell depletion) and 0.5×10^6^ transformed FTE cells were injected into the ovarian bursa of 6-to 8-week-old female NSG mice^23^. IVIS imaging was conducted per our previously published protocol^23^. Tumor growth was monitored for 7 weeks at which time tumors were dissected out and sent for histology.

### Mouse cell depletion

Mouse cells were depleted from the xenograft tumor to enrich for human cells using Mouse Cell Depletion Kit (Miltenyi Biotec, 130-104-694) as previously described^24^. Briefly, tumor tissues were mechanically and enzymatically dissociated into single-cell suspension. After determining the number of isolated cells, cells were resuspended in 80 µl of buffer (1x PBS with 0.5% BSA) per 2 x 10^6^ tumor cells and 20 µl of the magnetic labeling reagent for mouse cells. Cells were then incubated at 4°C for 15 minutes. After incubation, sample volume was adjusted to 500 µl, using buffer, and run through LS columns for magnetic separation. Flow-through containing purified human cells was collected, and cells were then cultured and propagated using DMEM medium supplemented with 10% FBS and penicillin/streptomycin (penicillin: 100 units/mL, streptomycin: 0.1 mg/mL).

### DNA isolation

The isolated post-initiation and matched pre-initiation FTE were processed for DNA isolation using Qiagen DNeasy Blood and tissue kit (cat # 69504), following the manufacture’s protocol.

### Whole-genome sequencing

Sequence libraries were generated from isolated post-initiation and matched pre-initiation FTE using the Illumina WGS DNA Prep workflow (Illumina) and subsequently sequenced on the Illumina NovaSeq6000 platform with a minimum coverage of 50x and 30x, respectively. FASTQ files were filtered for quality, using FASTQC (v.0.11.8), and contaminants, using FastQScreen (v0.11.4), and trimmed of adapters, N-content, and low-quality bases (ea-utils (v1.04)). Quality FASTQ files containing paired-end clean reads were then mapped to the hg38 human reference genome with BWA (v0.7.17), and the resulting BAM files were sorted, merged by lane, and duplicate reads marked using Sambamba (v0.6.7) and Picard (v2.18.9) <u>(Tarasov et al.</u> <u>2015; Li et al. 2009)</u>. GATK BaseRecalibrator (v<u>4.4.0.0</u>) was used to adjust bases<u>(DePristo et al. 2011)</u>.

### SNP and short INDEL variant calling

GATK HaplotypeCaller (v4.4.0.0) was used on pre-initiation FTE bam files to call germline base substitutions and short INDEL variants which were quality filtered and annotated according to the GATK Best Practices Pipeline^25^. Filtered germline variant call format (VCF) files were used as the “panel of normals” for GATK Mutect2 (v4.4.0.0) somatic variant calling^26^.Two variant callers were employed to call somatic base substitutions and short indels, using the pre-initiation FTE samples as normal: GATK Mutect2 and Strelka2 (v2.9.10), both using default parameters in tumor-normal mode^27^. Variant calls were normalized and decomposed using vt(v0.58), merged using GATK CombineVariants, and left trimmed using vt^28^. Variants that passed both callers, had at least five supporting reads, and bidirectional read support were deemed variants of high confidence; these variants were functionally annotated using GATK Funcotator (v4.4.0.0) and matched with reported ClinVar variations^29^.All variants of interest were manually interrogated using the Integrative Genomics Viewer^30^.

### Structural variant and CNV detection

Structural variants (SVs) were called with Parliament2 (v0.1.11), utilizing the overlap of the callers lumpy (v0.2.13), CNVnator (v0.3.3), and Breakdancer (v1.4.3), genotyped with SVTyper (v0.7.0), and merged by SURVIVOR (v1.0.3), using default parameters.

### Mutational signature analysis

Mutational signature analysis was run on high-confidence variants, using the R package MutationalPatterns (v3.12.0)^31^. Variants were converted to categories of mutational spectra for single-base substitution (SBS) and double-base substitution (DBS). SBS and DBS catalogs were obtained from the COSMIC Mutational Signatures database (v99) and compared to sample mutational signatures by calculating the cosign similarity between corresponding count matrices^32^.

## Results

### hrMSCs are present in the microenvironment of STIC lesions

Our previous work demonstrated that cancer-educated CA-MSCs are present in virtually all invasive HGSOC cases^13^. However, the timing of CA-MSCs development remained unclear therefore we first asked whether similar cells are found surrounding pre-invasive STIC lesions. We previously demonstrated that the high expression of WT1 can distinguish CA-MSCs from normal MSCs^13^. To visualize and quantify CA-MSC-like cells in the STIC microenvironment, we developed a multi-spectral Vectra imaging panel to identify WT1-positive MSCs. All tissue samples were verified for histopathologic features of normal vs STIC-containing vs invasive HGSOC containing fallopian tubes by a board-certified gynecologic pathologist (Fig. 1A). Tissues were analyzed for CA-MSC localization and abundance by Vectra multispectral imaging (Fig. 1A-C). MSCs were identified in the stroma by co-expression of the classic MSC surface markers CD73, CD90, and CD105. We further distinguished CA-MSC-like cells from normal MSCs (nMSCs) by WT1 expression (CD45-/CD73+/CD90+/CD105+/WT1+). Given the lack of invasive carcinoma in our STIC samples, we herein refer to the CA-MSC-like cells (WT1+ MSCs) as high risk MSCs or hrMSCs. Intriguingly, we observed hrMSCs within the stroma adjacent to and underlying STIC lesions (Fig. 1A; red x), while hrMSCs were rarely detected in normal patient fallopian tubes.

**Figure 1.**
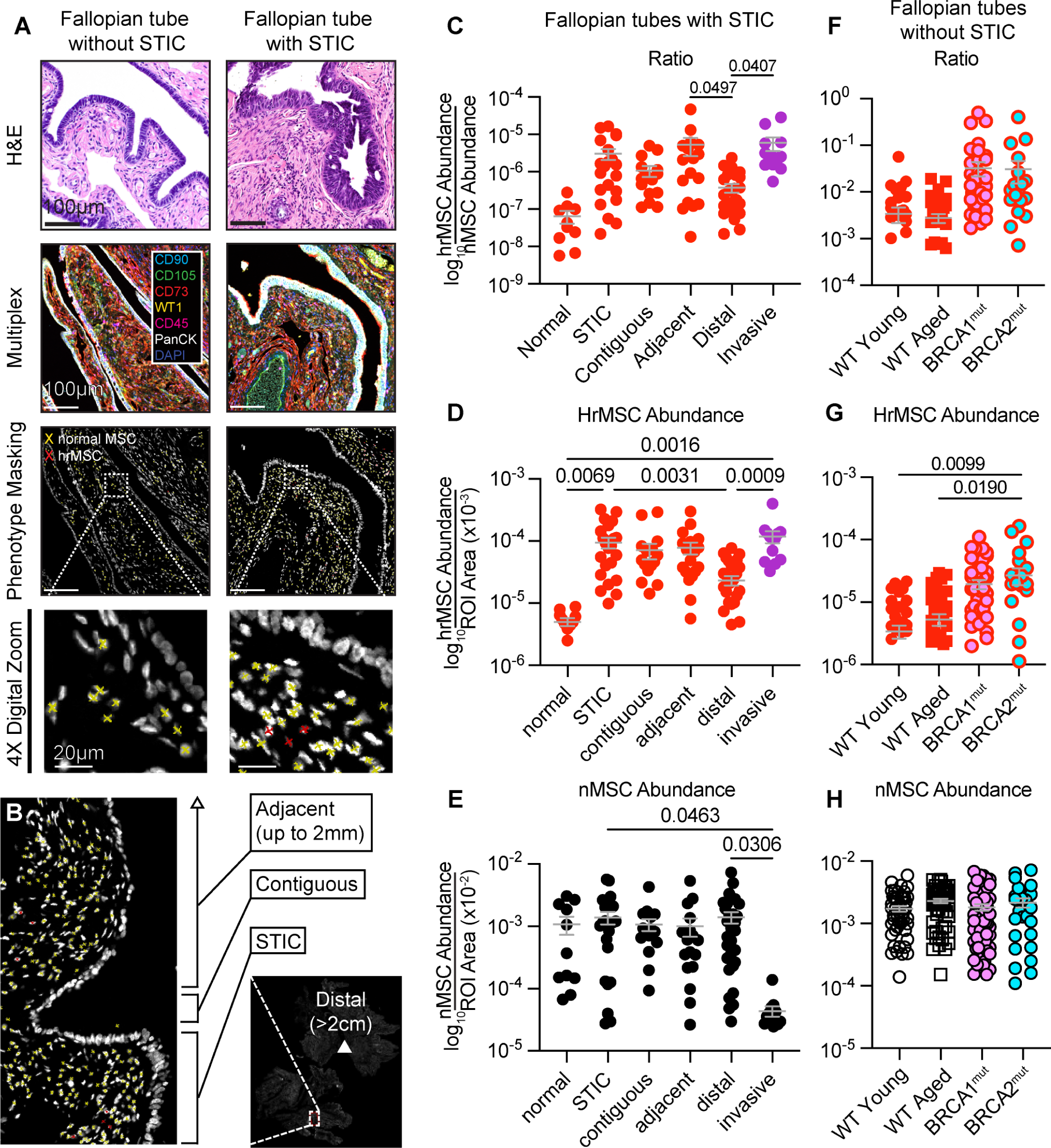
HrMSCS are enriched in the stroma underlying and adjacent to fallopian tube STIC lesions. A) Representative H&E-stained tissue sections of normal fallopian tubes (N=2) as well as fallopian tubes harboring STIC lesions (N=9) and/or HGSOC lesions (N=3). Histology is paired with multispectral immunofluorescence images of each respective group shown. Black and white images denote cells with the phenotypes of interest. Red x’s indicate hrMSCs or CA-MSCs (WT1+/CD73+/CD90+/CD105+/CD45-), respectively, while yellow x’s indicate nMSCs (WT1-/CD73+/CD90+/CD105+/CD45-). B) Schematic showing the criteria for our Vectra quantification. C) HrMSC to nMSC ratios. B) HrMSC abundance normalized to the area of our ROIs. E) NMSC Abundance normalized to the area of our ROIs. Individual data points are representative of an individual field. F) Ratio of hrMSCs to nMSCs in our Canary patient cohorts consisting of Wildtype (WT; N=10 young, N=9 aged), BRCA1 mutant (N=9), or BRCA2 mutant (N=6) healthy fallopian tubes lacking STIC/HGSOC. G) Quantification of hrMSC abundance and H) nMSC abundance are shown. P-values are shown in Fig. 1C-H. P-values were determined by ordinary one-way ANOVA with Tukey’s multiple comparisons analysis. Table summary of patient information/inclusion/exclusion criteria.

To further assess hrMSC spatial distribution and quantify the number of hrMSCs within the fallopian tube surrounding STIC lesions, we selected patient samples with STIC lesions that were incidental (N=6) and STIC lesions with concurrent invasive disease (N=11) and compared these to patient samples with invasive HGSOC within the fallopian tube (N=4) or completely normal fallopian tubes (N=2). We sought to determine if there was a stromal “field effect,” with hrMSCs involving stroma that extends beyond the boundaries of the STIC lesion. We thus quantified the abundance of hrMSCs and nMSCs directly underlying the STIC lesions as well as the stroma underlying FTE contiguous to the STIC region (first 20 normal, monolayered epithelial cells directly contiguous to the STIC lesion on both sides), the stroma underlying FTE adjacent to the STIC region (within 2mm of the STIC lesion on both sides), and the stroma underlying distal FTE (>2cm away from STIC lesions) (Fig. 1B-E). We compared these groups to hrMSCs found in normal fallopian tubes and CA-MSCs found within the invasive HGSOC microenvironment. Given that most STICs occur in the distal fimbriated end of the FT, we limited our investigation to this portion of the FT. Raw cell numbers (hrMSC or nMSC) were quantified and normalized by the tissue area analyzed and presented as hrMSC and nMSC abundance and hrMSC/nMSC ratio. Importantly, hrMSCs were significantly enriched in the stroma directly underlying STIC epithelium (Fig.1D). This was equivalent to CA-MSC abundance in invasive carcinoma. HrMSC abundance was also increased in contiguous and adjacent areas but decreased to significantly lower abundance in the stroma distal to STIC epithelium. Normal MSC abundance remained consistent across groups but was decreased in the stroma of invasive carcinoma (Fig. 1E). The ratio of hrMSC to nMSC was significantly higher in STIC-associated stroma and remained high in the contiguous and adjacent regions with a decrease in the regions distal to the STIC lesions. However, even in the distal regions, both the total number of hrMSCs and the ratio of hrMSC to nMSC remained higher than in normal FT (though this did not reach statistical significance). Interestingly, the abundance of hrMSCs found surrounding incidental STIC lesions (STIC lesions without associated invasive cancer) and STIC lesions with concurrent invasive cancer identified elsewhere were not statistically different (Supp. Fig. 1C).

We also demonstrated that hrMSCs are present in the fallopian tubes of women without STIC lesions (completely histologically normal tubes) (Fig. 1B). The initial two normal fallopian tubes analyzed were from women with a germline BRCA1 mutation. Given the 30-40 fold increased risk of developing HGSOC in patients with germline BRCA1 or BRCA2 mutations (*BRCA1^mut^ or BRCA2^mut^*), we hypothesized that hrMSCs are enriched in germline mutation carrier patient fallopian tubes which may account for some of the increased risk of developing HGSOC. We obtained additional normal FTs (without any identifiable pre-cancerous lesions or other pathology) from *BRCA1^mut^*carriers (N=9), *BRCA2^mut^* carriers (N=6), and *BRCA1/2*^wt^ patients (N=19). Given the increased risk of HGSOC with age, we divided the *BRCA*^wt^ patients into two age groups: 30-36 years old and 60-72 years old. The prevalence of hrMSCs was significantly higher in the *BRCA2^mut^* carriers, with a trend towards increased prevalence in the *BRCA1^mut^*carriers compared to *BRCA1/2^wt^* patients (Fig. 1F/G). There was also a trend toward increased hrMSC prevalence in the older *BRCA1/2*^wt^ patients compared to the younger patients (Fig. 1F/G), but this did not reach statistical significance.

### hrMSCs have the epigenomic and transcriptomic pattern of tumor-supportive CA-MSC**s**

We previously demonstrated that CA-MSCs have unique DNA methylation profiles that distinguish them from nMSCs^13,33^. Therefore we utilized the Infinium Methylation EPIC array to compared the DNA methylation pattern of MSCs from normal fallopian tubes of women with BRCA1/2*^mut^*to MSCs isolated from BRCA1/2*^wt^* normal fallopian tube, ovary or omental tissue or CA-MSCs isolated from invasive HGSOC (involving the fallopian tube, ovary or omentum) (Fig. 2A)^13^. As mentioned previously, patient fallopian tubes were analyzed by SEE-FIM (sectioning and extensively examining the fimbriated end) protocols^34^ to validate the absence of STIC or invasive cancer. Uniform Manifold Approximation and Projection (UMAP) and unsupervised hierarchical clustering revealed that roughly half of *BRCA1/2^mut^* fallopian tubes contained MSCs (red circles) which clustered with CA-MSCs (purple circles) while the rest of *BRCA1/2^mut^* MSCs clustered with normal MSCs (Fig. 2B). This is consistent with our Vectra imaging data demonstrating the presence of hrMSCs in some FTs of women with *BRCA1/2* mutations even in the absence of STIC or invasive cancer (Fig.1).

**Figure 2.**
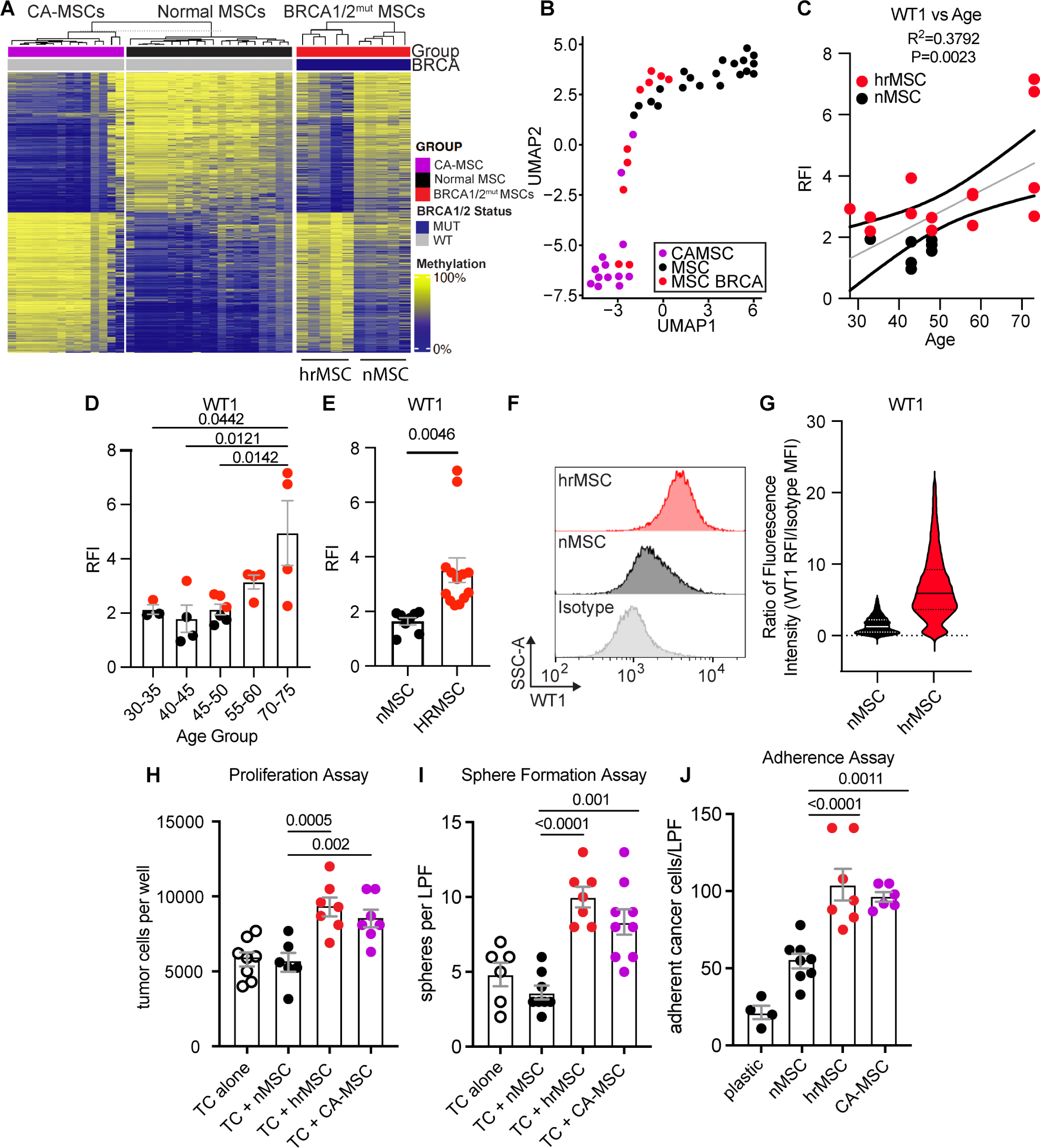
hrMSCs exhibit the epigenetic and phenotypic profile of CA-MSCs. A) DNA methylation array represented as a heat map. MSCs were taken from benign patient tissues (nMSCs; black circles; N=20), BRCA1/2^mut^ carrier patients without cancer (red circles; N=10), and patients with confirmed invasive HGSOC (CA-MSC; purple circles; N=14). B) UMAP on MSCs derived from patients in 2A. C) Relative fluorescence intensity (RFI) of WT1 405 was determined by flow cytometry for Wildtype (N=5), BRCA1^mut^ (N=3), and BRCA2^mut^ (N=13) MSCs. WT1 405 RFI was plotted against patient age. Linear regression of WT1 405 MFI versus patient age (gray line); R^2^=0.3745. D) MSCs were categorized into age groups. E) MSCs were classified into nMSCs or hrMSCs based on WT1 405 RFI with a minimum cut off of 2 RFI for hrMSCs. This categorization was independent of BRCA status and age. F/G) Representative histograms and violin plots of nMSC and hrMSC WT1 expression are shown to demonstrate heterogeneity of WT1 expression within a single patient line. Ovarian tumor cells (TC) were grown independently or in co-culture with nrMSCs, hrMSCs, or CA-MSCs H) under adherent, I) or non-adherent conditions. Individual cells and spheroids were counted at day 7 and graphed (n=3). J) Tumor cell adherence to nMSCs, hrMSCs, and CA-MSCs is shown (n=3). For Fig. 2D-J, p-values were determined by ordinary one-way ANOVA with Tukey’s multiple comparison analysis.

Given that the expression of WT1 is critical to CA-MSC development and can distinguish tumor supportive CA-MSCs from normal MSCs^13^, we next quantified WT1 protein expression across normal fallopian derived MSCs (from either BRCA1/2^wt^ or BRCA1/2^mut^) via flow cytometry (Fig. 2C-G). We used WT1 expression from CA-MSCs as our positive control. All CA-MSC cell lines express WT1 at ≥ 2 RFI (relative fluorescence intensity; Supp. Fig. 2B) while nMSCs fall below 2 RFI. We used this threshold of 2 to define a fallopian tube MSC as a hrMSC or nMSC (hrMSC= WT1 RFI ≥2, nMSC= WT1 RFI <2) (Fig. 2E). When WT1 expression was assessed in fallopian tube MSCs across the age spectrum we found that WT1 expression significantly increased with patient age and significantly more hrMSCs were present in the fallopian tubes of older women (Fig. 2C/D; R^2^=0.3792). The average age of women with hrMSCs was 10 years older than those with nMSCs (53-year-old vs 43-year-old). In some cases, WT1 expression varied between contralateral tubes of the same patient (Fig. 2F/G). It is important to note that MSCs derived from a patient were not uniformly WT1 high or WT1 low, but these populations existed on a spectrum (Fig. 2F) indicating a mixed population or a population with evolving WT1 expression (Fig. 2G). These data are consistent with our multi-spectral imaging data above demonstrating that hrMSCs are enriched in aged fallopian tubes and can be distinguished from nMSCs by WT1 expression.

### hrMSCs enhance cancer cell growth, sphere formation, and bind ovarian tumor cells

To determine if hrMSCs are phenotypically similar to their CA-MSC counterparts, i.e., have tumor-supportive function, we tested the impact of hrMSCs on critical aspects of HGSOC biology. When possible, assays were performed using matched hrMSCs and nMSCs (hrMSCs and nMSCs derived from contralateral fallopian tubes from the same woman to control for patient heterogeneity). We first tested the impact of hrMSCs on HGSOC proliferation. Cell Trace Violet (CTV) labeled HGSOC cells (OVCAR3, OVSAHO, and primary patient line 412) were grown with hrMSCs, nMSCs, or primary HGSOC-derived CA-MSCs at a 1:1 ratio and were counted over time. We utilized a 1:1 ratio in co-culture for the remainder of our experiments. HrMSCs increased the proliferation of HGSOC cells 2-fold compared to nMSCs (Fig. 2H). This was equivalent to the proliferation enhancement demonstrated by CA-MSC controls. We previously demonstrated that a critical function of CA-MSC support is their ability to directly bind cancer cells and form heterocellular complexes under non-adherent conditions^13^. We therefore tested the ability of hrMSCs to enhance tumor cell sphere formation. Sphere assays were performed with cancer cells grown with MSCs (nMSC, hrMSCs, or CA-MSCs) under non-adherent conditions, and the number of cancer cell-containing spheres were quantified. The number of cancer cell-containing spheres was equivalent between hrMSC and CA-MSC co-culture groups while nMSCs failed to enhance sphere formation (Fig 2I). We also compared the ability of hrMSCs to directly bind to cancer cells. MSCs (nMSCs, hrMSCs, or CA-MSCs) were grown in a monolayer, and fluorescently labeled cancer cells (OVCAR3, OVSAHO, or pt412) were tested for their ability to bind to MSCs. Cancer cells bound to hrMSCs at a level equivalent to CA-MSCs (∼3-to 4-fold higher than nMSCs) (Fig2J). These data demonstrate that hrMSCs are functionally equivalent to HGSOC-derived CA-MSCs in their support of ovarian cancer cell proliferation, sphere formation, and adherence to cancer cells.

### hrMSC tumor-supportive functions give rise to increased proliferation and increased DNA damage in FTE

Given the presence of hrMSCs within normal aging and BRCA1/2*^mut^*FTs and their abundance surrounding STIC lesions, we hypothesized that hrMSCs play a role in FTE transformation and HGSOC initiation. We first characterized the effects of hrMSCs on FTE cell growth and sphere formation. We conducted 2D proliferation assays and 3D sphere-formation assays under direct co-culture conditions. Primary patient-derived, germline *BRCA1/2^mut^* carrier FTE or immortalized FTE (FT190 control) cells were labelled with CTV and co-cultured with hrMSCs or nMSCs. FTE cells were counted daily, starting at day 0. By day 4, hrMSCs and nMSCs both significantly increased the growth of FTE compared to FTE grown alone (Fig. 3A). When co-cultured under non-adherent conditions, however, only hrMSCs significantly increased the growth of FTE spheroids while nMSCs did not (Fig 3B). CA-MSCs also enrich the stem-like population of tumor cells *in vitro*^13^. We therefore tested the ability of hrMSCs to impact the stemness of FTE. We quantified ALDH activity, which is a marker of stem cell capacity, in FTE co-cultured with hrMSCs or nMSCs. FTE cells that were co-cultured with hrMSCs demonstrated significantly increased ALDH activity compared to FTE cells cultured alone or in co-culture with nMSCs (Fig. 3C). This data indicates that hrMSCs also support the growth and potential stemness in primary, non-transformed epithelial cells.

**Figure 3.**
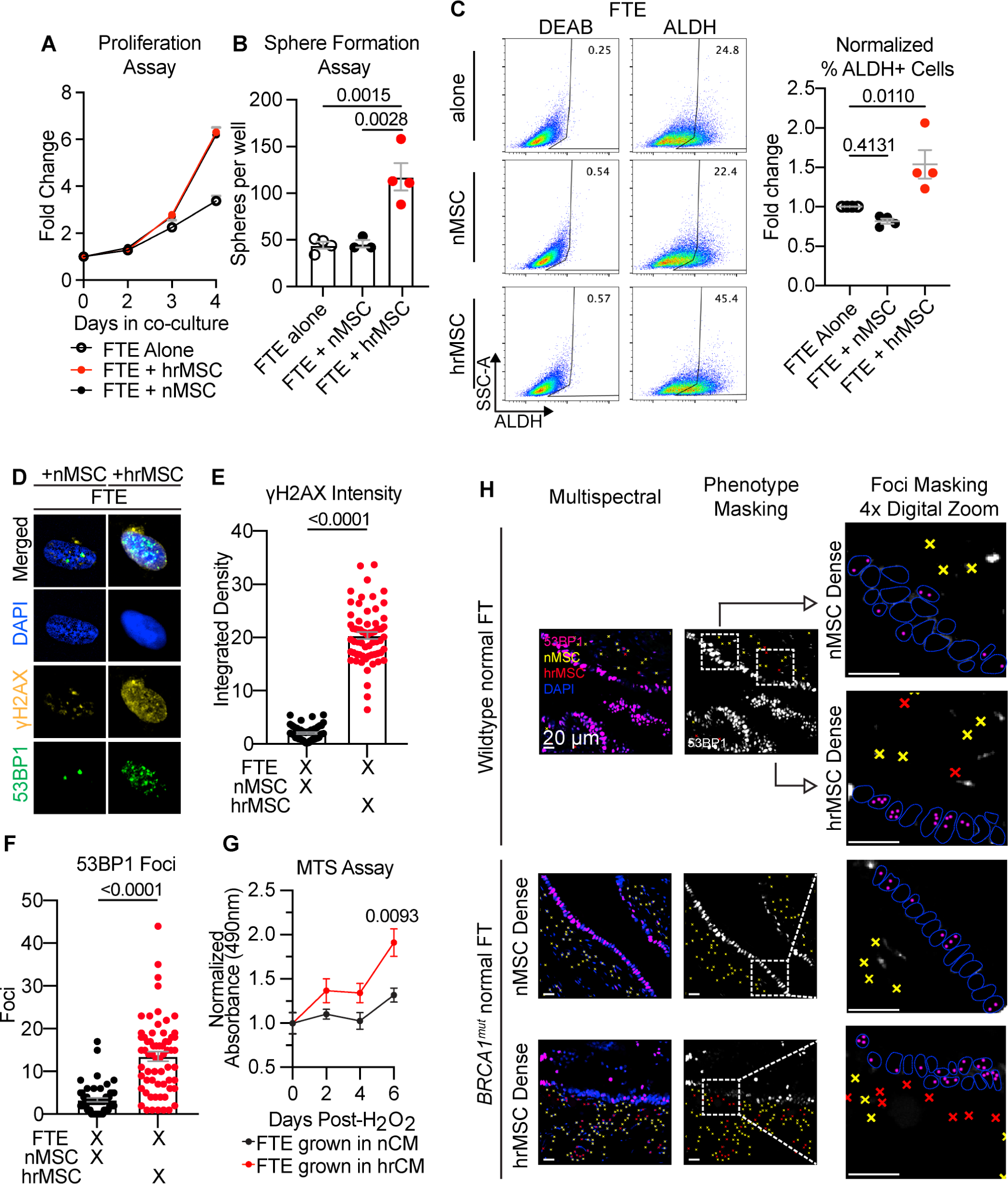
hrMSCs increase FTE proliferation, stemness, induce DNA damage in FTE, and enhance recovery of FTE following oxidative stress. A) Primary FTE were grown in co-culture with matched nMSCs, hrMSCs, or alone for up to 4 days. Each day the total number of viable cells were determined(n=3). B) Primary FTE were grown under non-adherent conditions with or without nrMSCs and hrMSCs. Spheroids were grown for 7 days and manually counted (n=3). C) Percentage of ALDH+ FT190 cells were determined by flow cytometry after 5 days of adherent co-culture with or without nrMSCs or hrMSCs. Percent of total cells that are ALDH+ are shown as well as representative flow plots (n=3). Primary FTE and immortalized FT190 control cells were co-cultured for 24 hours alone or with nMSCs or hrMSCs. Cells were assayed for E) γH2AX fluorescence intensity and F) 53BP1 foci per nucleus by fluorescent microscopy (n>3). Individual data points reflect individual nuclei. G) Primary FTE and FT190 control cells were treated with 50µM hydrogen peroxide and supplemented with either nMSC conditioned media (CM) or hrMSC CM. G) Cell viability was determined by MTS assay (n>3). H) Representative multispectral images of FTE with DNA DSBs (53BP1 foci) overlying either nMSC dense or hrMSCs dense stroma in WT or BRCA1^mut^ fallopian tubes. For Fig. 3A-G, p-values were determined by one-way ANOVA with Tukey’s multiple comparison analysis.

DNA damage, DNA double strand breaks, and DNA repair defects are well known hallmarks of HGSOC, therefore, we next assessed the impact of hrMSCs on FTE DNA damage. Primary FTE was co-cultured with matched hrMSCs or nMSCs. We assayed FTE for generalized DNA damage with γH2AX and DNA double-strand breaks (DSB) by 53BP1 foci. 24-hours after co-culture, FTE that were co-cultured with hrMSCs exhibited robust DNA damage and DNA DSBs (Fig. 3D-F). Importantly, we validated this finding in primary patient samples in situ. We applied our hrMSC Vectra multi-spectral imaging panel and incorporated staining for 53BP1 foci in normal BRCA1/2^wt^ and BRCA1^mut^ fallopian tubes (without any pathologic findings). We observed that in areas with high prevalence of hrMSCs, the overlying FTE demonstrated a larger relative abundance of 53BP1 foci compared to FTE overlying nMSC enriched regions (Fig. 3H).

The high burden of DNA DSBs in FTE exposed to hrMSCs led us to question if hrMSCs confer a survival and/or recovery advantage to FTE following DNA damage. To test this, we treated FTE with 50 µM of hydrogen peroxide for 10 minutes in FBS-free media. This dose is the minimum effective dose used to elicit cell death in the FTE cells. Following treatment, we neutralized the peroxide with media containing FBS and supplemented FTE with conditioned media from nMSCs or hrMSCs every other day for 6 days. At days 2, 4, and 6 post-treatment, viable FTE cells were quantified via MTS assay. Indeed, hrMSC conditioned media promotes increased cell recovery following peroxide treatment relative to nMSC controls (Fig. 3G). These data lead us to conclude that, not only are hrMSCs promoting FTE cell growth and stemness, they are also inducing DNA damage in the form of DNA DSBs and supporting FTE recovery following DNA damage.

### hrMSCs induce full malignant transformation

Given the striking impact of hrMSCs on FTE DNA damage, recovery, and proliferation, we next tested the impact of hrMSCs on FTE malignant transformation. We first created organoids from primary patient-derived, benign FTE with (i) *p53*-null mutation (*p53^mut^* FTE) or (ii) germline heterozygous *BRCA1^mut^* (BRCA1*^mut^* FTE) with hrMSCs or nMSC in a 1:1 FTE:MSC ratio. FTE-alone or MSC-alone organoids were also created as controls. Organoids developed within 7 days and were maintained in culture for 10 weeks. Representative *p53^mut^* FTE + nMSC or hrMSC organoids are shown in Supp. Fig. 4A. Organoids were then dissociated to ensure efficient injection and prevent mechanical shearing. Single cell suspensions were normalized based on epithelial cell number—1×10^5^ FTE per injection and injected into the mammary fat of NSG mice. Organoids containing *p53^mut^* FTE or *BRCA1^mut^* FTE alone failed to initiate tumors. Organoids with hrMSCs alone or nMSCs alone also did not initiate tumors. Similarly, organoids containing *p53^mut^* FTE or *BRCA1^mut^* FTE grown with nMSCs did not initiate tumors over 12 months. Strikingly, *p53*^mut^ FTE + hrMSC organoids (2 of 3) and BRCA1^mut^ FTE + hrMSCs (1 of 3) initiated tumors within 4 months (Fig. 4A-C). The two *p53*^mut^ + hrMSC organoids which initiated tumors also developed metastatic disease to the lung and liver. Representative images of the primary tumors and metastatic tumors are shown (Fig. 4B).

**Figure 4.**
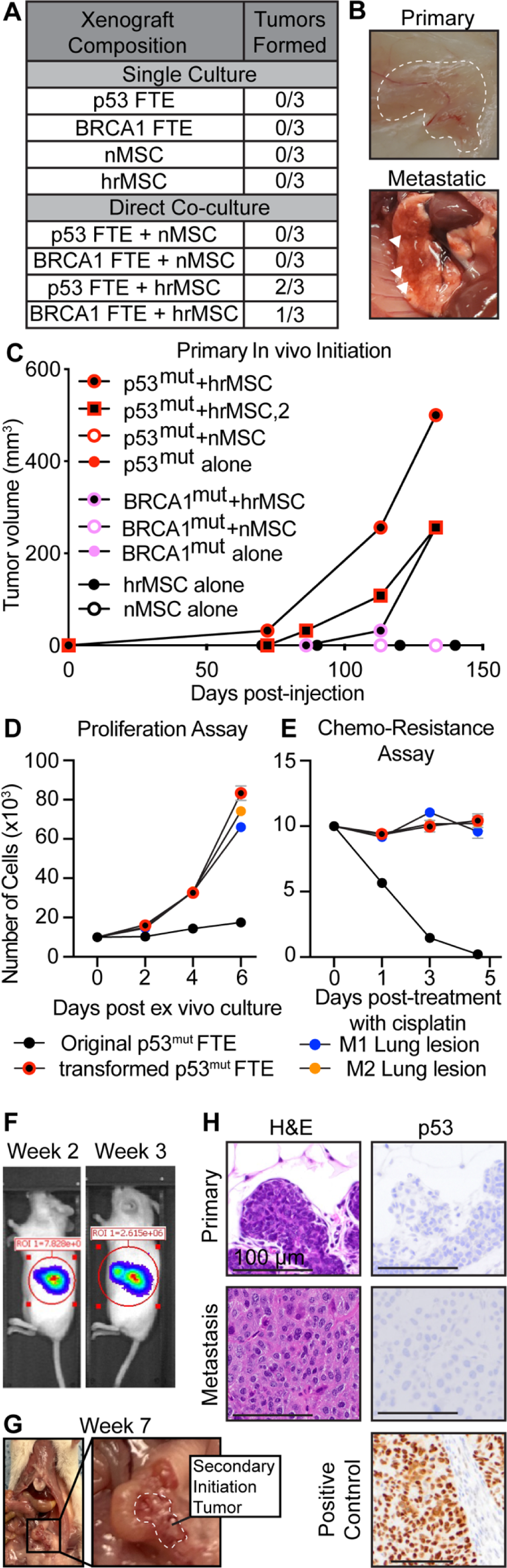
hrMSCs induce malignant transformation of p53^mut^ and BRCA1^mut^ primary FTE cells. Primary FTE p53 mutant cells were cultured under non-adherent conditions alone, or with either nMSCs or hrMSCs. Co-cultures were grown for 10 weeks then dissociated and injected in NSG mice. A) Total number of mice that developed tumors for each condition is shown (n=1). B) Macroscopic representative images of primary tumors and secondary metastases. C) Tumor volume per mouse was graphed over the course of 5 months. D) Tumors cells from the primary and metastatic tumors were excised, dissociated, and passaged up to 6 days. E) Tumor cells were also treated with 1 mg/mL cisplatin and cell growth was determined up to 5 days (n=3). F) IVIS imaging of a representative mouse after week 2 and week 3 of secondary initiation (n=1). G) Gross anatomy of tumor initiation site. Tumors were allowed to grow for 2 weeks before IVIS imaging began. H) Tumors were excised and processed for H&E. Canonical histologic features of HGSOC are shown.

We next assessed functional changes in transformed FTE which initiated tumors. *P53*^mut^ FTE were isolated from primary tumors and lung lesions 6 months post-injection. Human cells were enriched from the xenografts, using mouse cell depletion kits. Residual human MSCs or differentiated human stroma were removed with FACs sorting on the EPCAM+, CD73/90/105-cell population. The transformed *p53*^mut^ FTE that we isolated were propagated *ex vivo* and were plated for proliferation and chemoresistance assays. Compared to the original, primary *p53*^mut^ FTE, the transformed *p53*^mut^ FTE demonstrated a significantly higher proliferation rate and increased resistance to cisplatin treatment (Fig. 4D/E). Additionally, the transformed *p53*^mut^ FTE demonstrated immortalization without replicative senescence. Compared to the parent *p53*^mut^ FTE line, in which cells underwent senescence after 8-12 passages, the transformed *p53*^mut^ FTE have been passaged over 80 times with a doubling time of ∼24hrs. To further confirm full malignant transformation, we performed a secondary initiation experiment using an orthotopic model. The transformed *p53*^mut^ FTE, after *ex vivo* propagation, were injected into the ovarian bursa of NSG mice and allowed to grow for 3 months. To quantify the time to engraftment, a subset of the transformed *p53*^mut^ FTE were transduced with luciferase lentivirus to enable tracking with IVIS imaging. In this subset of tumors engraftment was seen by Week 2 (Fig. 4F) with evidence of metastasis by week 7 post-injection (Fig. 4G). All other mice bearing transformed *p53*^mut^ FTE (without luciferase to prevent any confounding of results from viral transfection) were sacrificed at 3 months to assess tumor initiation and burden. All mice (12 of 12) demonstrated tumor engraftment with local invasion. 5 of 12 mice demonstrated lung and liver metastasis. Tumors were excised and validated with histopathology (T.R.S.) demonstrating high grade carcinoma morphology, epithelial origin (Supp. Fig. 4B) and loss of p53 expression (consistent with the parent FTE line harboring a p53 null mutation) (Fig. 4H). Collectively, these data demonstrate a stromal-mediated FTE transformation.

### Transformed FTE exhibit genomic HGSOC mutation signatures

HGSOC bear hallmark mutation signatures and certain mutation signatures can infer potential mechanism of DNA damage ^7,35^ ^36^ ^37^. Thus, to characterize the genomic alterations in the transformed FTE and shed light on the underlying DNA damage mechanism in our model, we conducted whole genome sequencing on our metastatic *p53^mut^ FTE* transformed cell lines. Compared to the non-transformed parental FTE cell line, transformed FTE cells acquired somatic mutations in genes associated with HGSOC, namely *BRCA1* and *APC*, as well as structural variants (Fig. 5A/B). Moreover, we performed single-base signature analysis on our whole-genome sequencing data and compared this to COSMIC mutational signatures. Interestingly, we discovered that each sample exhibited single-base mutational signatures 1, 2 and 5, with signature 5 having the strongest relative frequency (Fig. 5C). Signature 5 has unknown etiology; however, it has been associated with DNA damage resulting from replication stress, chronic exposure to reactive oxygen species (ROS), and aging. Based on this, we sought to characterize levels of oxidative stress in nMSCs and hrMSCs. We relied on CellROX oxidation probes to measure oxidative stress. We first observed that CellROX Deep Red (DR) positively correlated with WT1 expression in MSCs (Fig. 5E). Both CellROX DR and CellROX Green fluorescence intensities were significantly greater in hrMSCs compared to nMSC controls by flow cytometry (Fig. 5E) and immunofluorescence (IF) (Fig. 5G). Further, hrMSC oxidative stress was reduced to nMSC levels with a single 24-hour treatment of the antioxidant Trolox (Fig. 5F).

**Figure 5.**
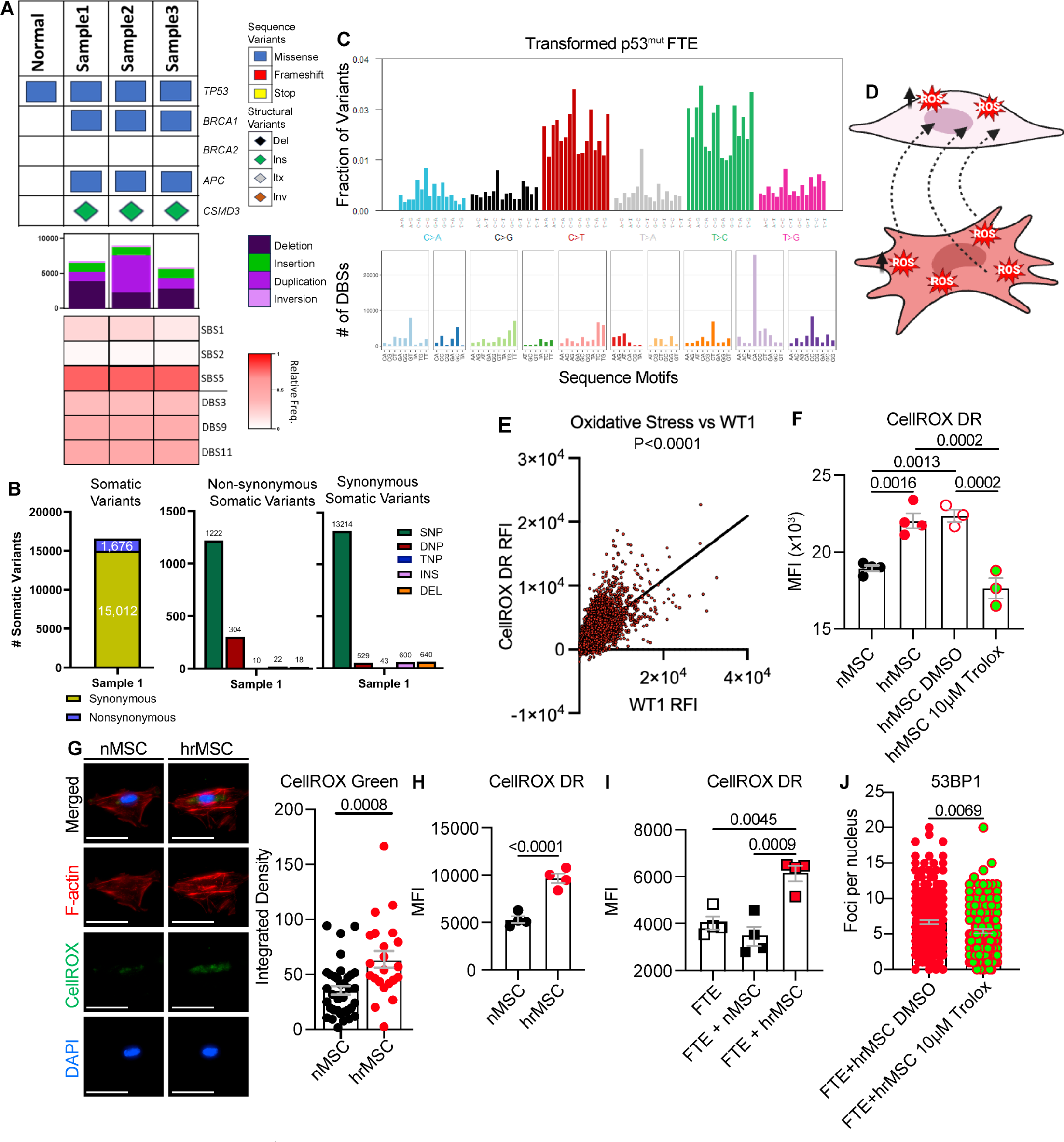
Transformed p53^mut^ FTE that metastasized harbor mutational hallmarks of HGSOC. Whole genome sequencing was used to characterize mutations present in metastatic samples from the primary initiation of transformed p53^mut^ FTE. A) Oncoprint including mutations in genes commonly mutated in HGSOC. Correlations to COSMIC single base and double base signatures are shown (SBS and DBS, respectively). B) Summary statistics of mutational analysis including synonymous, and non-synonymous mutations. C) Representative graph of single base mutations for one sample exhibiting the pan-cancer mutational signature (signature 5). Representative graph of the non-specific double base signature. D) We hypothesize that hrMSCs have have greater oxidative stress that is communicated to FTE to induce DNA damage. E) Linear regression of WT1 relative fluorescence intensity (RFI) and the oxidation probe CellROX Deep Red (CellROX DR) determined by flow cytometry. Individual data points represent individual events. F) CellROX DR staining was compared between nMSCs and hrMSCs by flow cytometry. 10µM Trolox in DMSO was used as a negative control. Individual data points represent individual wells (n>3). G) CellROX Green fluorescence intensity was compared between nMSCs and hrMSCs by fluorescent microscopy. Representative cells are shown at 20X magnification with a 100µm scale bar. Individual data points represent individual cells. >3 fields per group were taken. Student’s T-test was used to determine significance. H/I) FTE were co-cultured with either nMSCs or hrMSCs for 24 hours and assayed for CellROX DR stain by flow cytometry (n=3). Representative cells are shown at 20X magnification with a 100µm scale bar. Individual data points represent individual nuclei. >3 fields were taken per condition. J) FTE were co-cultured with hrMSCs ±10µM Trolox in DMSO for 24 hours. 53BP1 foci per nucleus were determined by fluorescence microscropy. For figures 5E-H, p-values were determined by ordinary one-way ANOVA with Tukey’s multiple comparison analysis.

Given that we observed DNA damage and mutation in FTE exposed to hrMSCs, we next tested if FTE co-cultured with hrMSCs also acquired oxidative stress. Therefore, we labeled hrMSCs with CTV and FTE with CFSE. Cells were co-cultured for 24-hours and were harvested for analysis by flow. We observed significantly greater CellROX DR MFI in FTE co-cultured with hrMSCs relative to FTE co-cultured alone or with nMSCs (Fig. 5H/I). nMSCs elicited a non-significant shift in FTE oxidative stress.

We next tested if hrMSC oxidative stress was essential for DNA damage in FTE. We co-cultured hrMSCs with FTE ±5 µM Trolox, and quantified 53BP1 foci. We observed a significant reduction in 53BP1 foci in FTE cells that were co-cultured with hrMSCs in the presence of Trolox (Fig. 5J). These data indicate that hrMSCs are inducing DNA damage in FTE through oxidative stress by either direct or indirect mechanisms.

### hrMSCs have increased oxidation at baseline and produce elevated lipid hydroperoxides relative to matched, non-tumor-supportive nMSCs

One consequence of increased oxidative stress is lipid peroxidation. As lipid peroxides are more stable than ROS, can be trafficked between cells, and can break down into the highly mutagenic compounds malondialdehyde (MDA) and 4-hydroxy-2E-nonenal (4-HNE) we hypothesized this may be an important mechanism of hrMSC induced DNA damage in FTE. We characterized 4-HNE and MDA (Fig. 6A-D) levels in nMSCs and hrMSCs via IF and flow cytometry, respectively. Both 4-HNE and MDA were increased in hrMSCs relative to nMSCs. Trolox treatment decreased levels of both 4-HNE and MDA. Like CellROX, both 4-HNE and MDA positively correlated with WT1 expression (Fig. 6E,F). To understand if lipid peroxides are being trafficked from hrMSCs to FTE cells, we utilized both ELISA and IF assays to detect extracellular and intracellular MDA. While MDA concentrations were near undetectable in the conditioned media of either nMSC or hrMSCs (Supp. Fig. 6A), MDA puncta are visualized by IF on the cell membrane of hrMSCs (Fig. 6H). Notably, these MDA puncta are present in/on the filopodia/nanotubes originating from CTV labelled hrMSCs and connect to FTE suggesting that transfer of lipid peroxides occured via direct interaction. In support of this, co-culture of FTE with hrMSCs also resulted in significantly increased MDA and 4-HNE in FTE compared to FTE alone or FTE co-cultured with nMSCs (Fig. 6I,J). We next verified that exogenous lipid peroxide breakdown products could induce DNA DSBs in FTE. We treated FTE with 20µM 4-HNE for 8 hours and quantified 53BP1 foci. Indeed, 53BP1 foci were significantly increased in FTE treated with 4-HNE compared to FTE treated with DMSO (Fig. 6K). These data indicates that hrMSCs induce DNA DSBs in FTE, in part via production and trafficking of lipid peroxide break down products.

**Figure 6.**
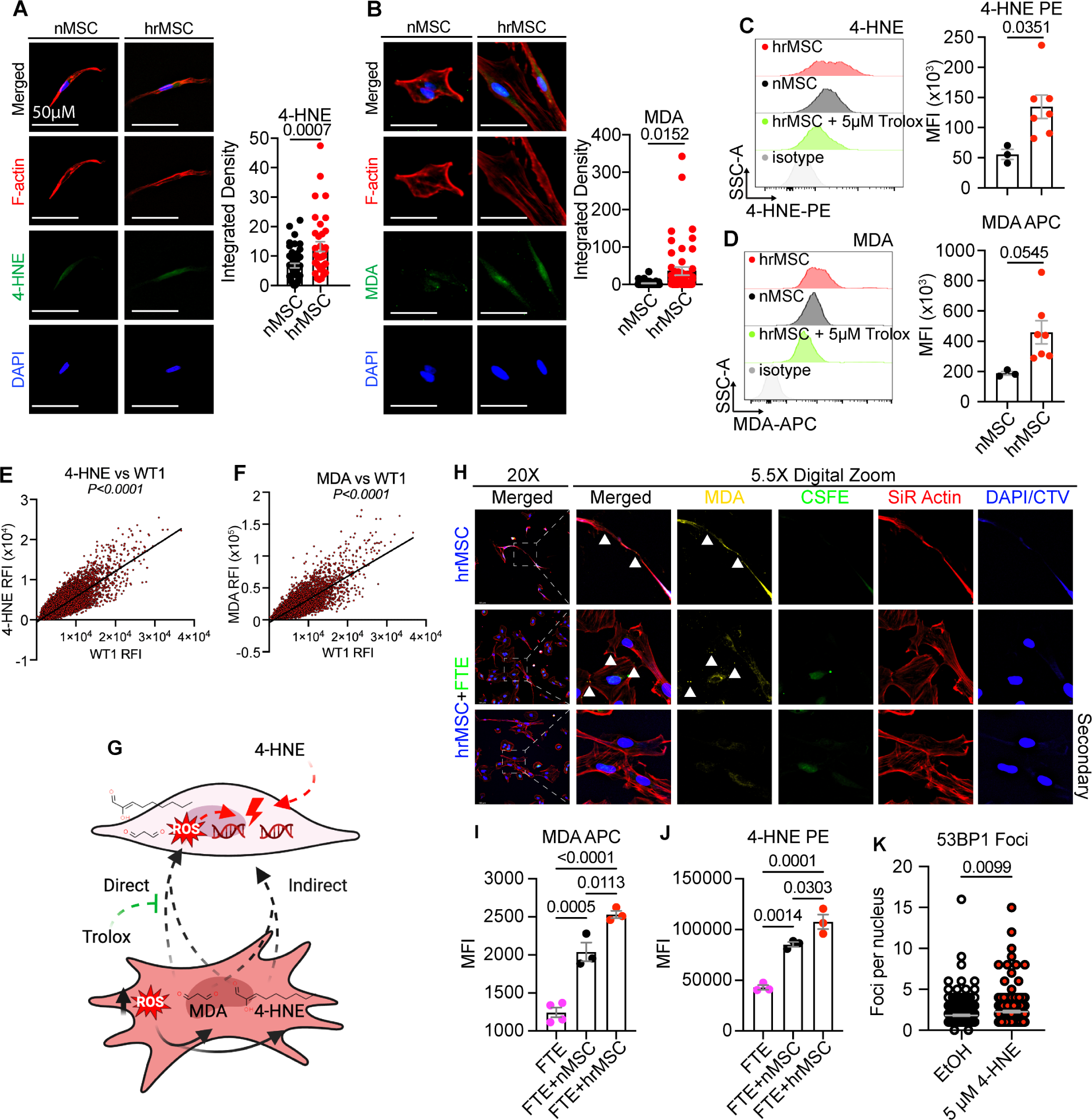
HrMSCs exhibit increased oxidative stress and lipid peroxide breakdown products that induce DNA damage in FTE. 4-hydroxy-2E-nonenal (4-HNE) and malondialdehyde (MDA) were compared between nMSCs and hrMSCs by A/B) fluorescence microscopy (n=2) and C/D) flow cytometry (n>3). A/B) Each data point is an individual cell with N=3 nMSC lines and N=7 hrMSC lines. 20X magnification representative images are shown. >3 fields were imaged and analyzed per condition. C/D) Each data point is the MFI determined for an individual patient cell line. 10µM Trolox treatment is included as a negative control. Each point is a separate patient.E/F) Linear regression between E) 4-HNE and WT1 and F) MDA and WT1 by flow cytometry. Individual points are independent events. G) We hypothesize that FTE DNA damage is a result of hrMSC lipid peroxidation. Dashed lines are test in the following figure panels. H) Representative images of MDA in hrMSCs (CTV+) that are co-cultured with FTE (CT CSFE+). Arrows depict MDA puncta that are visible within nanotubule connections between cells (n>3). Quantification of I) 4-HNE-PE and J) MDA-APC fluorescence intensities in FTE cells and MSCs with or without co-culture. K) FTE were treated with 5µM 4-HNE and assayed for 53BP1 foci. P-values were determined by ordinary one-way ANOVA with Tukey’s multiple comparison analysis.

### Restoration of pAMPKα1/AMPKα1 in hrMSCs ameliorates FTE DNA damage burden after co-culture

Next, we sought to clarify the mechanism by which hrMSCs develop increased oxidative stress and the DNA damage inducing phenotype. Increased oxidative stress in the setting of reduced AMP-associated protein kinase alpha 1 (AMPKα1) activation and feedback inhibition is a well-characterized hallmark of aging^38,39^. Therefore, we compared AMPKα1 and phospho-AMPKα1 (T172; pAMPKα1) protein expression between nMSCs and hrMSCs from multiple patients. While the pAMPKα1:AMPKα1 ratio remains unchanged, nMSCs express significantly higher AMPKα1 and pAMPKα1 compared to hrMSCs (Fig. 7A). When we restored AMPKα1 and pAMPKα1 expression pharmacologically using BC1618 (Fig. 7B), a small molecule inhibitor of pAMPKα1 degradation, we observed dose dependent decreases in total hrMSC MDA and 4-HNE via flow cytometry (Fig. 7C). These data suggest that reduced AMPKα1 expression may contribute to hrMSC DNA damage function (Fig. 7D). Therefore, we induced expression of AMPKα1 via lentiviral transduction of hrMSCs with GFP-tagged AMPKα1. We flow sorted GFP+ hrMSCs and confirmed overexpression via western blot (Fig. 7E). We observed reduced MDA/4HNE levels in hrMSCs overexpressing AMPKα1 compared to hrMSCs transduced with empty vector controls (Fig. 7F). Similarly, FTE co-cultured with hrMSCs overexpressing AMPKα1 exhibited reduced MDA/4-HNE compared to FTE cells co-cultured with empty vector controls (Fig. 7G). Lastly, when FTE cells were co-cultured with AMPKα1 overexpressing hrMSCs, total 53BP1 foci were reduced as well as the number of cells exhibiting increased 53BP1 foci (Fig. 7H). Collectively, this data demonstrated AMPKα1 expression is reduced in hrMSCs resulting in reduced feedback inhibition of oxidative stress/lipid peroxidation and subsequent induction of DNA damage in co-cultured FTE.

**Figure 7.**
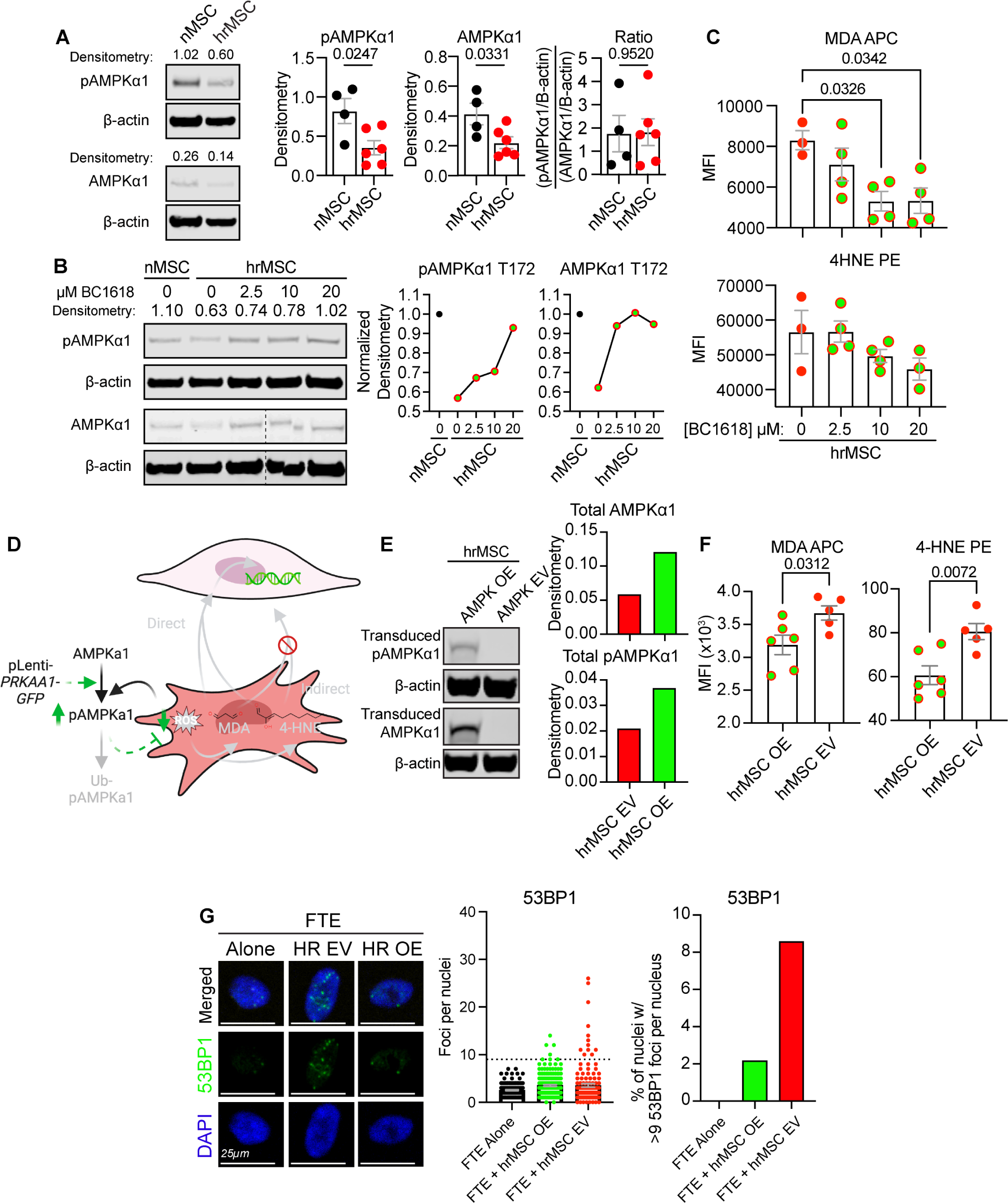
Pharmacologic and genetic AMPKα1 restoration results in reduced LPBPs and DNA damage in FTE. A) Western blot and quantification of AMPKα1 and pAMPKα1 from nMSC (N=4) and hrMSCs (N=6) patient lines. B) Western blot and quantification of AMPKα1 and pAMPKα1 following BC1618 treatment for 24 hours. C) BC1618 treated hrMSCs were analyzed by flow cytometry for 4-HNE-PE and MDA-APC fluorescence intensity. D) We hypothesize that restoration of hrMSC AMPKα1 expression will ameliorate DNA damage in FTE cells. E) Western blot and quantification of MSCs transduced with lentiviral particles containing either GFP-tagged AMPKα1 (hrMSC OE) or GFP-empty vector constructs (hrMSC EV). Endogenous and exogenous AMPKα1 and pAMPKα1 were combined for quantification. F) Quantification of 4-HNE-PE and MDA-APC fluorescence intensity by flow cytometry in transduced cell lines (n=2). G) Representative 20X images of FTE that were co-cultured at a 1:1 ratio with transduced hrMSCs. 53BP1 foci were quantified by fluorescence microscopy. >3 fields per condition were imaged and analyzed. P-values were determined by ordinary one-way ANOVA with Tukey’s multiple comparison analysis.

## Discussion

It is now recognized that the majority of HGSOC is primarily derived from the fallopian tube epithelium; therefore, the majority of studies focus exclusively on FTE intrinsic selective pressures for oncogenesis ^7,8,40–42^. Further, pathologic mutations in either *BRCA1* or *BRCA2* convey a 30-to 40-fold increased risk of developing HGSOC; therefore, most studies focus on patients with germline mutations in these genes. However, the majority of HGSOC occurs in women without germline *BRCA1*/2 mutations, highlighting a critical need to elucidate a mechanism of ovarian cancer initiation that considers other, more broadly applicable factors ^1^.

Our work indicates a critical role for the fallopian tube stroma in ovarian cancer initiation. Using a combination of multispectral immunofluorescence, DNA methylation analysis, and flow cytometry for WT1 expression, we identified a subset of MSCs that exist within the fallopian tube stroma of women without cancer which phenocopy fully cancer-educated CA-MSCs. CA-MSCs are characterized by their tumor-supportive functions; however, given that these MSCs were identified in women without cancer, we termed these cells high-risk MSCs or hrMSCs. We demonstrated that hrMSCs exist within the fallopian tube stroma of women without HGSOC. HrMSCs were most abundant surrounding STIC lesions which are considered the precursor lesion to HGSOC. Additionally, the presence of hrMSCs extended well beyond the boundaries of the STIC lesion, creating a possible field effect surrounding and contributing to the development of STIC lesions and, ultimately, HGSOC. HrMSCs were also found in women without any identifiable pathology in their fallopian tubes and were enriched in patients with greater age and patients with germline *BRCA1*/2^mut^; however, they were not exclusive to those subgroups. We also demonstrated that hrMSCs functionally mirror CA-MSCs in their ability to support tumor cell growth. Moreover, hrMSCs also directly influenced benign FTE. hrMSCs increased DNA damage and DSB burden in FTE. This association was validated *in vivo*, with co-localization of FTE with 53BP1 foci (a marker of DNA DSBs) with hrMSCs within the underlying stroma. DNA DSBs are considered hallmarks of HGSOC, often directly contributing to the allelic copy number variations typical of invasive disease. Given the toxicity of DNA DSBs in human cells, we expected total loss of epithelium after prolonged co-culture or upon additional cell stress. Therefore, it was striking to discover that not only do FTE proliferate more when in co-culture with hrMSCs, but FTE recover significantly better, after additional oxidant stress, when exposed to hrMSC conditioned media. These data indicate that hrMSCs provide two vital stimuli for ovarian cancer initiation: induction of DNA damage and promotion of epithelial survival. Additionally, hrMSCs may support the stem-like capacity of FTE by enriching cells with high ALDH expression and with the capacity to grow in spheres. As many hypothesize that cancer initiation depends on malignant transformation of the normal stem cell population, the impact of stromal-mediated DNA damage induction and survival may be even more potent when occurring in the FTE stem cell population^43^.

Most importantly, hrMSCs triggered full malignant transformation of primary FTE *in vivo*. These findings were validated by WGS, which demonstrated classic hallmarks of HGSOC. To date, the transformed epithelium has been passaged >80 times in comparison to primary epithelial lines that senesce within weeks of *ex vivo* culture, demonstrating permanent retention of malignant features resulting from hrMSC influence. Other sources of FTE DNA damage have been described; however, none have recapitulated *in vivo* transformation. These additional stressors may encourage the pro-tumorigenic function of hrMSCs; therefore, future investigation into hrMSC metabolism, cell-to-cell interactions, and FTE responses are vital in determining the precise mechanism of epithelial transformation.

Our initial investigation into the mechanism of hrMSC-mediated malignant transformation was based on the genomic analysis of the transformed FTE, which demonstrated a mutational signature associated with chronic oxidative stress (COSMIC single base signature 5). We then verified that hrMSCs have higher oxidative stress compared to normal MSCs. This high oxidative stress leads to an increased burden of 4-HNE and MDA in hrMSCs. 4-HNE and MDA belong to two separate lipid peroxide breakdown product families, each capable of inducing genotoxic stress, either in the form of replication fork stalling or DNA DSBs. Lipid peroxides have been implicated in disease progression; however, their involvement in HGSOC initiation has been neglected. hrMSCs express low levels of AMPKα1, which is known to result in reduced responsivity toward fluctuations in cellular oxidative stress. This suggests that hrMSCs lack robust, oxidant stress feedback inhibition, which would permit hrMSCs to accumulate 4-HNE/MDA and induce DNA damage in FTE. Intriguingly, pharmacologic, and genetic restoration of AMPKα1 in hrMSCs or hrMSC treatment with Trolox resulted in significantly lower oxidative stress, 4-HNE/MDA burden, and FTE DNA damage and DSBs. Further work is needed to understand the mechanism driving lower AMPKα1 in hrMSCs; how hrMSCs both induce DNA damage yet increase FTE survival; and other related or orthogonal mechanisms underlying hrMSC-mediated FTE transformation.

Collectively, this work identifies the stromal microenvironment as a crucial factor in HGSOC initiation. We demonstrate that benign resident MSCs take on a pro-tumorigenic phenotype prior to the initiation of cancer and that these cells actively participate in the malignant transformation of overlying epithelial cells. This information is vitally important to our understanding of how HGSOC forms and in the effort to create early detection or prevention strategies for this deadly disease. To date, biomarkers associated with HGSOC have not reached the necessary sensitivity and specificity to be useful for early detection of ovarian cancer. Understanding the role of the fallopian tube stroma in HGSOC initiation may enable the discovery of stroma-based biomarkers to improve the development of methods for early detection. Similarly, stroma-based prevention strategies may also be possible to block the tumor-promoting impact of hrMSCs. This work highlights the importance of the stromal microenvironment in HGSOC initiation and provides support to investigate similar stroma-mediated functions at other sites of oncogenesis.

## Acknowledgements

We would like to thank ProMark and the Pathology Biospecimen Core for tissue acquisition. We would like to thank the imaging, flow cytometry and animal core facilities. The following sources contributed funding for this work: The Honorable Tina Brozman Foundation (LC, RD) DoD Omics Consortium W81XWH-22-1-0852 (LC, RB, RD), DOD Ovarian Cancer Research Program IIRA OC210139 (LC), NIH HCC Ovarian Cancer SPORE P50CA272218-01A1 (LC, RB), NIH U01 AG077923 (LC, RB, TF), NIH P50 CA228991 SPORE in ovarian cancer (RD), The Miriam and Sheldon G. Adelson Medical Research Foundation (RD).

## References

1. Siegel, R.L., Miller, K.D., Wagle, N.S., and Jemal, A. (2023). Cancer statistics, 2023. CA Cancer J. Clin. 73, 17–48. 10.3322/caac.21763.

2. Burke, W., Barkley, J., Barrows, E., Brooks, R., Gecsi, K., Huber-Keener, K., Jeudy, M., Mei, S., O’Hara, J.S., and Chelmow, D. (2023). Executive summary of the ovarian cancer evidence review conference. Obstet. Gynecol. 142, 179–195. 10.1097/AOG.0000000000005211.

3. Mei, J., Tian, H., Huang, H.-S., Hsu, C.-F., Liou, Y., Wu, N., Zhang, W., and Chu, T.-Y. (2021). Cellular models of development of ovarian high-grade serous carcinoma: A review of cell of origin and mechanisms of carcinogenesis. Cell Prolif. 54, e13029. 10.1111/cpr.13029.

4. Reavis, H.D., and Drapkin, R. (2020). The tubal epigenome - An emerging target for ovarian cancer. Pharmacol. Ther. 210, 107524. 10.1016/j.pharmthera.2020.107524.

5. Shih, I.-M., Wang, Y., and Wang, T.-L. (2021). The origin of ovarian cancer species and precancerous landscape. Am. J. Pathol. 191, 26–39. 10.1016/j.ajpath.2020.09.006.

6. Zhang, S., Dolgalev, I., Zhang, T., Ran, H., Levine, D.A., and Neel, B.G. (2019). Both fallopian tube and ovarian surface epithelium are cells-of-origin for high-grade serous ovarian carcinoma. Nat. Commun. 10, 5367. 10.1038/s41467-019-13116-2.

7. Labidi-Galy, S.I., Papp, E., Hallberg, D., Niknafs, N., Adleff, V., Noe, M., Bhattacharya, R., Novak, M., Jones, S., Phallen, J., et al. (2017). High grade serous ovarian carcinomas originate in the fallopian tube. Nat. Commun. 8, 1093. 10.1038/s41467-017-00962-1.

8. Perets, R., Wyant, G.A., Muto, K.W., Bijron, J.G., Poole, B.B., Chin, K.T., Chen, J.Y.H., Ohman, A.W., Stepule, C.D., Kwak, S., et al. (2013). Transformation of the fallopian tube secretory epithelium leads to high-grade serous ovarian cancer in Brca;Tp53;Pten models. Cancer Cell 24, 751–765. 10.1016/j.ccr.2013.10.013.

9. Bronder, D., Tighe, A., Wangsa, D., Zong, D., Meyer, T.J., Wardenaar, R., Minshall, P., Hirsch, D., Heselmeyer-Haddad, K., Nelson, L., et al. (2021). TP53 loss initiates chromosomal instability in fallopian tube epithelial cells. Dis. Model. Mech. 14. 10.1242/dmm.049001.

10. Zong, Y., Huang, J., Sankarasharma, D., Morikawa, T., Fukayama, M., Epstein, J.I., Chada, K.K., and Witte, O.N. (2012). Stromal epigenetic dysregulation is sufficient to initiate mouse prostate cancer via paracrine Wnt signaling. Proc Natl Acad Sci USA 109, E3395–404. 10.1073/pnas.1217982109.

11. Wu, J., Raz, Y., Recouvreux, M.S., Diniz, M.A., Lester, J., Karlan, B.Y., Walts, A.E., Gertych, A., and Orsulic, S. (2022). Focal Serous Tubal Intra-Epithelial Carcinoma Lesions Are Associated With Global Changes in the Fallopian Tube Epithelia and Stroma. Front. Oncol. 12, 853755. 10.3389/fonc.2022.853755.

12. Lochhead, P., Chan, A.T., Nishihara, R., Fuchs, C.S., Beck, A.H., Giovannucci, E., and Ogino, S. (2015). Etiologic field effect: reappraisal of the field effect concept in cancer predisposition and progression. Mod. Pathol. 28, 14–29. 10.1038/modpathol.2014.81.

13. Fan, H., Atiya, H.I., Wang, Y., Pisanic, T.R., Wang, T.-H., Shih, I.-M., Foy, K.K., Frisbie, L., Buckanovich, R.J., Chomiak, A.A., et al. (2020). Epigenomic Reprogramming toward Mesenchymal-Epithelial Transition in Ovarian-Cancer-Associated Mesenchymal Stem Cells Drives Metastasis. Cell Rep. 33, 108473. 10.1016/j.celrep.2020.108473.

14. Dominici, M., Le Blanc, K., Mueller, I., Slaper-Cortenbach, I., Marini, F., Krause, D., Deans, R., Keating, A., Prockop, D., and Horwitz, E. (2006). Minimal criteria for defining multipotent mesenchymal stromal cells. The International Society for Cellular Therapy position statement. Cytotherapy 8, 315–317. 10.1080/14653240600855905.

15. Karst, A.M., and Drapkin, R. (2012). Primary culture and immortalization of human fallopian tube secretory epithelial cells. Nat. Protoc. 7, 1755–1764. 10.1038/nprot.2012.097.

16. Zhou, W., Triche, T.J., Laird, P.W., and Shen, H. (2018). SeSAMe: reducing artifactual detection of DNA methylation by Infinium BeadChips in genomic deletions. Nucleic Acids Res. 46, e123. 10.1093/nar/gky691.

17. McInnes, L., Healy, J., and Melville, J. (2018). UMAP: Uniform Manifold Approximation and Projection for Dimension Reduction. arXiv. 10.48550/arxiv.1802.03426.

18. Barbolina, M.V., Adley, B.P., Shea, L.D., and Stack, M.S. (2008). Wilms tumor gene protein 1 is associated with ovarian cancer metastasis and modulates cell invasion. Cancer 112, 1632–1641. 10.1002/cncr.23341.

19. Hylander, B., Repasky, E., Shrikant, P., Intengan, M., Beck, A., Driscoll, D., Singhal, P., Lele, S., and Odunsi, K. (2006). Expression of Wilms tumor gene (WT1) in epithelial ovarian cancer. Gynecol. Oncol. 101, 12–17. 10.1016/j.ygyno.2005.09.052.

20. Taube, E.T., Denkert, C., Sehouli, J., Kunze, C.A., Dietel, M., Braicu, I., Letsch, A., and Darb-Esfahani, S. (2016). Wilms tumor protein 1 (WT1)--not only a diagnostic but also a prognostic marker in high-grade serous ovarian carcinoma. Gynecol. Oncol. 140, 494–502. 10.1016/j.ygyno.2015.12.018.

21. Yamamoto, S., Tsuda, H., Kita, T., Maekawa, K., Fujii, K., Kudoh, K., Furuya, K., Tamai, S., Inazawa, J., and Matsubara, O. (2007). Clinicopathological significance of WT1 expression in ovarian cancer: a possible accelerator of tumor progression in serous adenocarcinoma. Virchows Arch. 451, 27–35. 10.1007/s00428-007-0433-4.

22. Han, Y., Song, C., Zhang, T., Zhou, Q., Zhang, X., Wang, J., Xu, B., Zhang, X., Liu, X., and Ying, X. (2020). Wilms’ tumor 1 (WT1) promotes ovarian cancer progression by regulating E-cadherin and ERK1/2 signaling. Cell Cycle 19, 2662–2675. 10.1080/15384101.2020.1817666.

23. Atiya, H.I., Orellana, T.J., Wield, A., Frisbie, L., and Coffman, L.G. (2021). An Orthotopic Mouse Model of Ovarian Cancer using Human Stroma to Promote Metastasis. J. Vis. Exp. 10.3791/62382.

24. Agorku, D.J., Tomiuk, S., Klingner, K., Wild, S., Rüberg, S., Zatrieb, L., Bosio, A., Schueler, J., and Hardt, O. (2016). Depletion of Mouse Cells from Human Tumor Xenografts Significantly Improves Downstream Analysis of Target Cells. J. Vis. Exp. 10.3791/54259.

25. Van der Auwera, G.A., Carneiro, M.O., Hartl, C., Poplin, R., Del Angel, G., Levy-Moonshine, A., Jordan, T., Shakir, K., Roazen, D., Thibault, J., et al. (2013). From FastQ data to high confidence variant calls: the Genome Analysis Toolkit best practices pipeline. Curr. Protoc. Bioinformatics 11, 11.10.1-11.10.33. 10.1002/0471250953.bi1110s43.

26. Benjamin, D.I., Sato, T., Cibulskis, K., Getz, G., Stewart, C., and Lichtenstein, L. (2019). Calling Somatic SNVs and Indels with Mutect2. BioRxiv. 10.1101/861054.

27. Kim, S., Scheffler, K., Halpern, A.L., Bekritsky, M.A., Noh, E., Källberg, M., Chen, X., Kim, Y., Beyter, D., Krusche, P., et al. (2018). Strelka2: fast and accurate calling of germline and somatic variants. Nat. Methods 15, 591–594. 10.1038/s41592-018-0051-x.

28. Tan, A., Abecasis, G.R., and Kang, H.M. (2015). Unified representation of genetic variants. Bioinformatics 31, 2202–2204. 10.1093/bioinformatics/btv112.

29. Landrum, M.J., Lee, J.M., Benson, M., Brown, G.R., Chao, C., Chitipiralla, S., Gu, B., Hart, J., Hoffman, D., Jang, W., et al. (2018). ClinVar: improving access to variant interpretations and supporting evidence. Nucleic Acids Res. 46, D1062–D1067. 10.1093/nar/gkx1153.

30. Robinson, J.T., Thorvaldsdóttir, H., Winckler, W., Guttman, M., Lander, E.S., Getz, G., and Mesirov, J.P. (2011). Integrative genomics viewer. Nat. Biotechnol. 29, 24–26. 10.1038/nbt.1754.

31. Blokzijl, F., Janssen, R., van Boxtel, R., and Cuppen, E. (2018). MutationalPatterns: comprehensive genome-wide analysis of mutational processes. Genome Med. 10, 33. 10.1186/s13073-018-0539-0.

32. Tate, J.G., Bamford, S., Jubb, H.C., Sondka, Z., Beare, D.M., Bindal, N., Boutselakis, H., Cole, C.G., Creatore, C., Dawson, E., et al. (2019). COSMIC: the catalogue of somatic mutations in cancer. Nucleic Acids Res. 47, D941–D947. 10.1093/nar/gky1015.

33. Coffman, L.G., Pearson, A.T., Frisbie, L.G., Freeman, Z., Christie, E., Bowtell, D.D., and Buckanovich, R.J. (2019). Ovarian Carcinoma-Associated Mesenchymal Stem Cells Arise from Tissue-Specific Normal Stroma. Stem Cells 37, 257–269. 10.1002/stem.2932.

34. Medeiros, F., Muto, M.G., Lee, Y., Elvin, J.A., Callahan, M.J., Feltmate, C., Garber, J.E., Cramer, D.W., and Crum, C.P. (2006). The tubal fimbria is a preferred site for early adenocarcinoma in women with familial ovarian cancer syndrome. Am. J. Surg. Pathol. 30, 230–236. 10.1097/01.pas.0000180854.28831.77.

35. Steele, C.D., Abbasi, A., Islam, S.M.A., Bowes, A.L., Khandekar, A., Haase, K., Hames-Fathi, S., Ajayi, D., Verfaillie, A., Dhami, P., et al. (2022). Signatures of copy number alterations in human cancer. Nature 606, 984–991. 10.1038/s41586-022-04738-6.

36. Alexandrov, L.B., Nik-Zainal, S., Wedge, D.C., Aparicio, S.A.J.R., Behjati, S., Biankin, A.V., Bignell, G.R., Bolli, N., Borg, A., Børresen-Dale, A.-L., et al. (2013). Signatures of mutational processes in human cancer. Nature 500, 415–421. 10.1038/nature12477.

37. Alexandrov, L.B., Nik-Zainal, S., Wedge, D.C., Campbell, P.J., and Stratton, M.R. (2013). Deciphering signatures of mutational processes operative in human cancer. Cell Rep. 3, 246–259. 10.1016/j.celrep.2012.12.008.

38. Stancu, A.L. (2015). AMPK activation can delay aging. Discoveries (Craiova) 3, e53. 10.15190/d.2015.45.

39. Salminen, A., and Kaarniranta, K. (2012). AMP-activated protein kinase (AMPK) controls the aging process via an integrated signaling network. Ageing Res. Rev. 11, 230–241. 10.1016/j.arr.2011.12.005.

40. Folkins, A.K., Jarboe, E.A., Saleemuddin, A., Lee, Y., Callahan, M.J., Drapkin, R., Garber, J.E., Muto, M.G., Tworoger, S., and Crum, C.P. (2008). A candidate precursor to pelvic serous cancer (p53 signature) and its prevalence in ovaries and fallopian tubes from women with BRCA mutations. Gynecol. Oncol. 109, 168–173. 10.1016/j.ygyno.2008.01.012.

41. Levanon, K., Ng, V., Piao, H.Y., Zhang, Y., Chang, M.C., Roh, M.H., Kindelberger, D.W., Hirsch, M.S., Crum, C.P., Marto, J.A., et al. (2010). Primary ex vivo cultures of human fallopian tube epithelium as a model for serous ovarian carcinogenesis. Oncogene 29, 1103–1113. 10.1038/onc.2009.402.

42. Karst, A.M., and Drapkin, R. (2010). Ovarian cancer pathogenesis: a model in evolution. J. Oncol. 2010, 932371. 10.1155/2010/932371.

43. Sell, S. (2010). On the stem cell origin of cancer. Am. J. Pathol. 176, 2584–2494. 10.2353/ajpath.2010.091064.

