## Supplementary material for "Stromal mediated DNA damage promotes high grade serous ovarian cancer initiation": Supplimental files

#### ***Supplemental Figures***

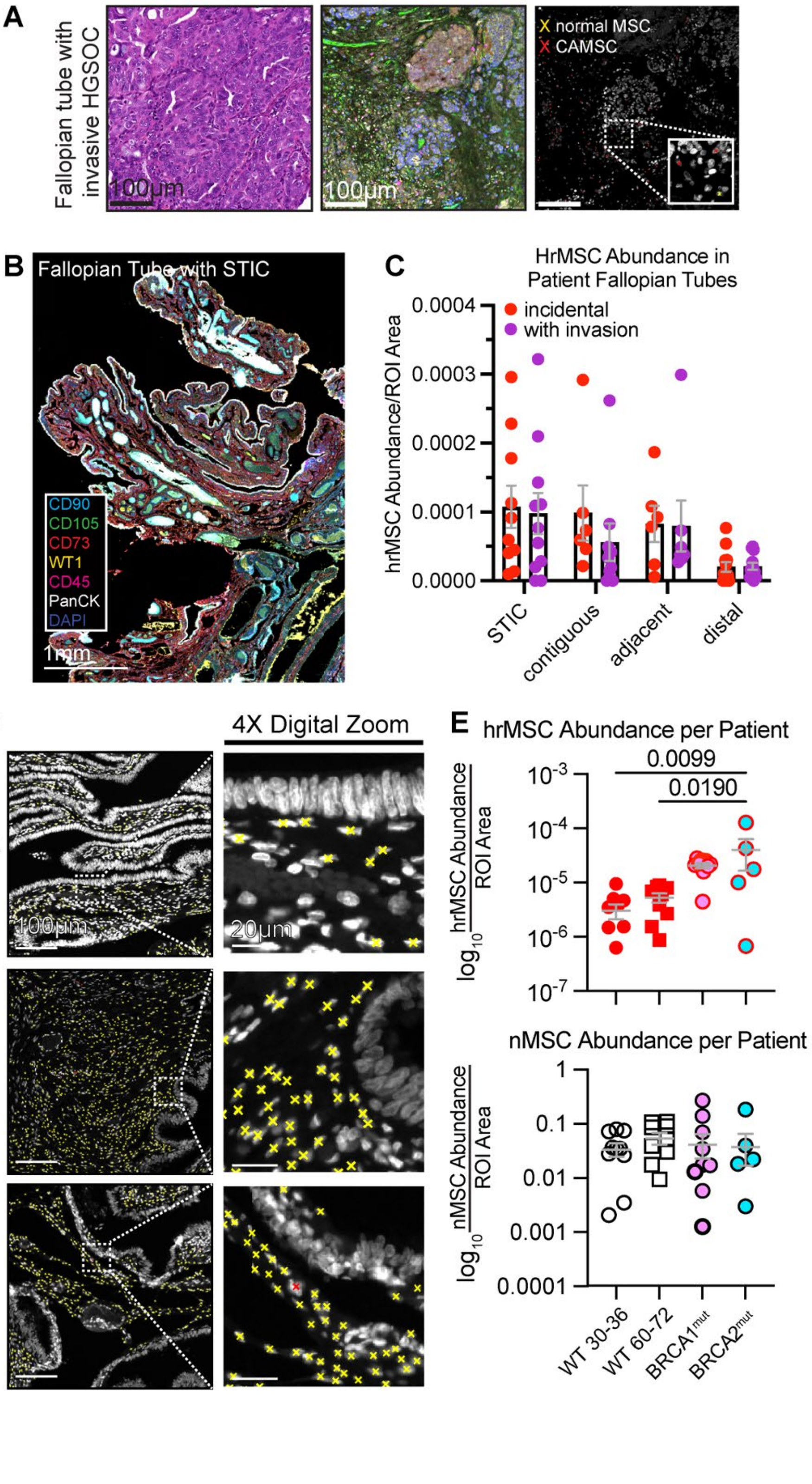


***Supp. Figure 1*** *A) Histology, Vectra, and MSC masking of HGSOC samples. B) Stitched, multiplexed imaged of a representative fallopian tube for reference. C) HrMSC abundance underlying incidental STIC or STIC accompanied by HGSOC. D) Representative phenotype masking of WT, BRCA1, and BRCA2 mutant fallopian tubes. E) hrMSC abundance and nMSC abundance per patient. P-values are shown and were determined by ordinary one-way ANOVA with Tukey's multiple comparisons analysis. P-values are reflected in Fig. 1G/H.*

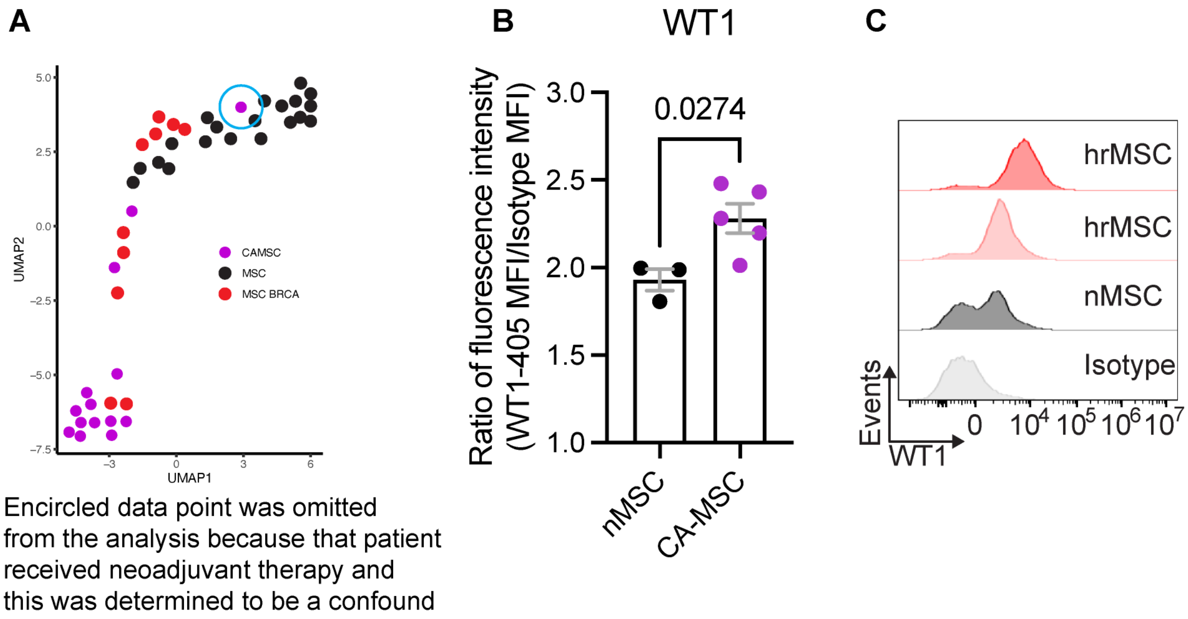


***Supp. Figure 2*** *A) Unmodified principal component analysis. B) WT1 MFI between CA-MSCs and nMSCs. The lower limit of CA-MSC WT1 MFI was used to classify hrMSCs. C) Representative histograms demonstrating heterogeneity of WT1 expression in patient cell lines. P-values are shown. P-values were determined by Student's T-test.*

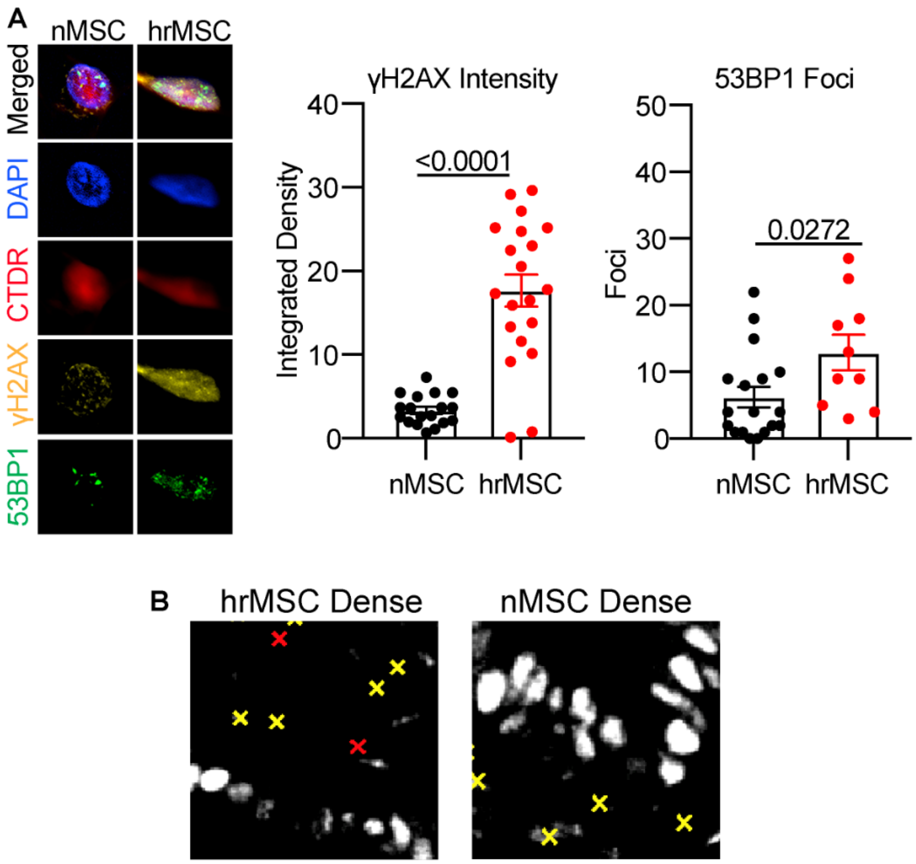


***Supp. Figure 3*** *A) gH2AX and 53BP1 foci in Cell trace deep red labelled MSCs. MSCs were co-cultured with FTE at a 1:1 ratio. B) Representative phenotype masking and 53BP1 stain in patient fallopian tubes. Foci were identified using ImageJ and are indicated as magenta foci in Fig. 3H. Cells that lacked clear foci, including cells without signal, were considered to be negative for 53BP1 foci. P-values are shown. P-values were determined by Student's T-test.*

Supplemental Data:

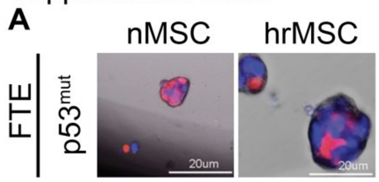


***Supp. Figure 4*** *A) Representative images of organoids that were injected into NSG mice. FTE and MSCs were seeded at a 1:1 ratio under non-adherent conditions. FTE are labelled in red while hrMSCs are labelled in blue.*

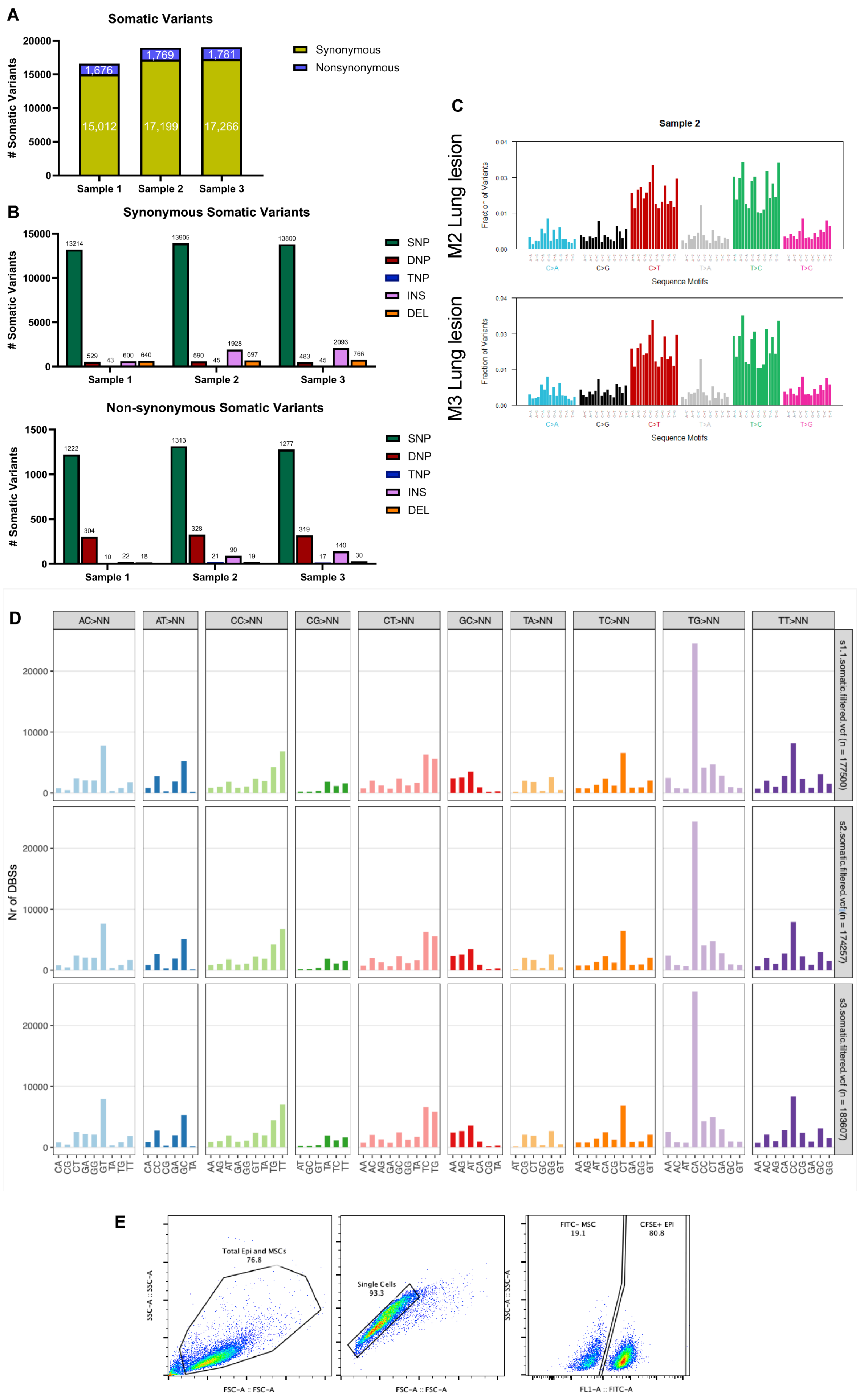
A
Somatic Variants
### Somatic Variants
20000
15000
10000
5000
0
1,676
15,012
1,769
17,199
1,781
17,266
Sample 1
Sample 2
Sample 3
Synonymous
Nonsynonymous
B
Synonymous Somatic Variants
### Somatic Variants
15000
10000
5000
0
13214
529
43
600
640
13905
590
45
1928
697
13800
483
45
2093
766
Sample 1
Sample 2
Sample 3
SNP
DNP
TNP
INS
DEL
Non-synonymous Somatic Variants
### Somatic Variants
1500
1000
500
0
1222
304
10
22
18
1313
328
21
90
19
1277
319
17
140
30
Sample 1
Sample 2
Sample 3
SNP
DNP
TNP
INS
DEL
C
Sample 2
M2 Lung lesion
Fraction of Variants
0.04
0.03
0.01
0.00
C>A
C>G
C>T
T>A
T>C
T>G
Sequence Motifs
M3 Lung lesion
Fraction of Variants
0.04
0.03
0.01
0.00
C>A
C>G
C>T
T>A
T>C
T>G
Sequence Motifs
D
AC>NN
AT>NN
CC>NN
CG>NN
CT>NN
GC>NN
TA>NN
TC>NN
TG>NN
TT>NN
Nr of DBSs
20000
10000
0
s1.1.somatic.filtered.vcf (n = 177500)
s2.somatic.filtered.vcf (n = 174257)
s3.somatic.filtered.vcf (n = 183607)
CA CG CT GA GG GT TA TG TT
CA CC CG GA GC TA
AA AG AT GA GG GT TA TC TG
AT GC GT TA TC TT
AA AC AG GA GC GG TA TC TG
AA AG AT CA CG TA
AT CG CT GC GG GT
AA AG AT CA CG CT GA GG GT
AA AC AT CA CC CT GA GC GT
AA AC AG CA CC CG GA GC GG
E
Total Epi and MSCs
76.8
SSC-A :: SSC-A
FSC-A :: FSC-A
Single Cells
93.3
FSC-A :: FSC-A
FITC- MSC
19.1
CFSE+ EPI
80.8
SSC-A :: SSC-A
FL1-A :: FITC-A

***Supp. Figure 5** A/B) Summary statistics for all three samples including further subdivided into synonymous and non-synonymous somatic variants. C) Single base mutation signatures in mouse 2 and mouse 3. D) Double base mutation signatures for all three mice that were sequenced. E) Representative gating strategy for co-culture flow experiments.*

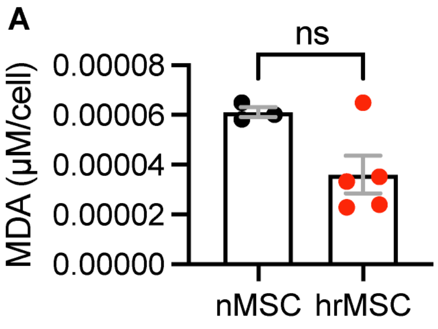


***Supp. Figure 6** A) MDA adduct ELISA comparing nMSC and hrMSC conditioned media. Individual data points represent individual patient cell lines. Statistics were determined by Student's T-test.*

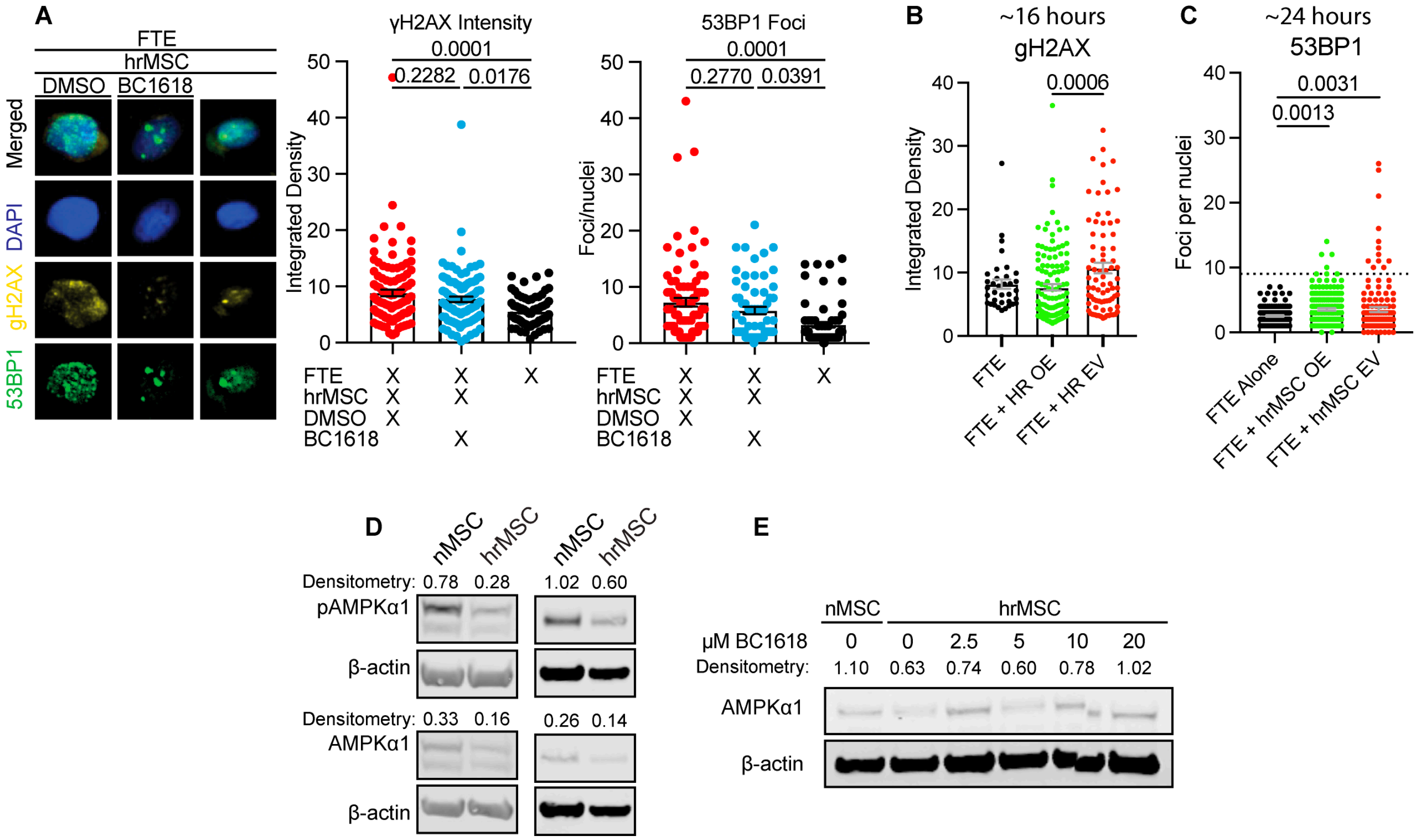


***Supp. Figure 7** A) hrMSCs were treated with 10 μM BC1618 for 24 hours then co-cultured with FTE at a 1:1 ratio for 24-hours. gH2AX integrated density and 53BP1 foci were quantified and are shown. B/C) FTE were co-cultured with hrMSCs overexpressing AMPKα1-mGFP or mGFP empty vector. B) gH2AX was quantified 16-hours post-seeding and C) 53BP1 foci were quantified at 24-hours post-seeding. D) Additional western blots from patient cell lines that were used in the AMPKα1 quantification. E) Unmodified western blot of hrMSCs AMPKα1 used in Fig. 7B. P-values are shown for Fig. A/B/C. P-values were determined by ordinary one-way ANOVA with Tukey's multiple comparisons analysis.*
